# A Temporal Interactome Atlas across RNA Viruses reveals Convergent Vulnerabilities for Antiviral Repurposing

**DOI:** 10.64898/2026.09.21.753288

**Authors:** Tavis. J. Reed, Alicja Tadych, Olga G. Troyanskaya, Ileana M. Cristea

**Affiliations:** Lewis-Sigler Institute for Integrative Genomics, Princeton University, Carl Icahn Laboratory, Princeton, NJ, 08544; Princeton Precision Health, Princeton University, 252 Nassau Street, Princeton, NJ, 08540; Department of Computer Science, Princeton University, 35 Olden Street, Princeton, NJ, 08540; Flatiron Institute, Simons Foundation, New York City, NY 10001, USA; Department of Molecular Biology, Princeton University, 119 Lewis Thomas Laboratory, Princeton, NJ, 08544

**Keywords:** Protein-protein interactions, proteomics, interaction networks, thermal proximity coaggregation, deep learning, drug repurposing, measles virus, HCoV-OC43, influenza A virus, dengue virus, zika virus

## Abstract

Viruses remodel host protein interactions to facilitate their replication, yet these dynamic interactomes remain largely unmapped and difficult to compare across infections. Here, to facilitate comparative virology studies, we build the Interactome and Viral Infection Dynamics (InterVir) Atlas, performing thermal proximity coaggregation to profile temporal interactome remodeling across five major RNA viral pathogens. We develop PROTEA, an AI framework that quantifies interactome homology and identifies convergent regulatory hubs. Infection-specific signatures include neuronal SHC1 rewiring associated with altered measles virus replication and dengue-specific secretase interactomes linked to cell-junction remodeling. Comparative virology analyses uncover unexpected interactome convergence between measles and flaviviruses, unobserved in protein-abundance profiling. Mapping these shared regulatory programs to drug targets allowed prioritization of compound repurposing as antivirals. Experimental testing of ten compounds demonstrated antiviral efficacy across multiple infections. Together, InterVir and PROTEA establish an open-access resource for comparative virology, facilitating the identification of conserved host vulnerabilities and drug repurposing.

## Introduction

Viruses are obligate intracellular parasites that, despite genetic simplicity, can have major impacts on human health. Human viruses exhibit large degrees of heterogeneity in genetic architecture, virion structure, tissue tropism, and disease pathogenesis^1–3^. Yet, all viruses share a fundamental requirement—they must systematically remodel their host cells to replicate^4–6^. Host remodeling can display shared features between different types of infections, such as virus-driven immune suppression, regulation of apoptosis, control of cellular metabolism, and the modulation of other key host pathways^6–13^. These finely tuned regulatory processes contribute to the impact of a virus on its micro-environment and compatibility for co-infections with other viruses. Critical for accomplishing either infection-specific or shared host remodeling is the ability acquired by viruses to temporally modulate host protein-protein interaction (PPI) networks that regulate a slew of critical cellular processes (Figure 1A)^11,12,14^.

**Figure 1.**
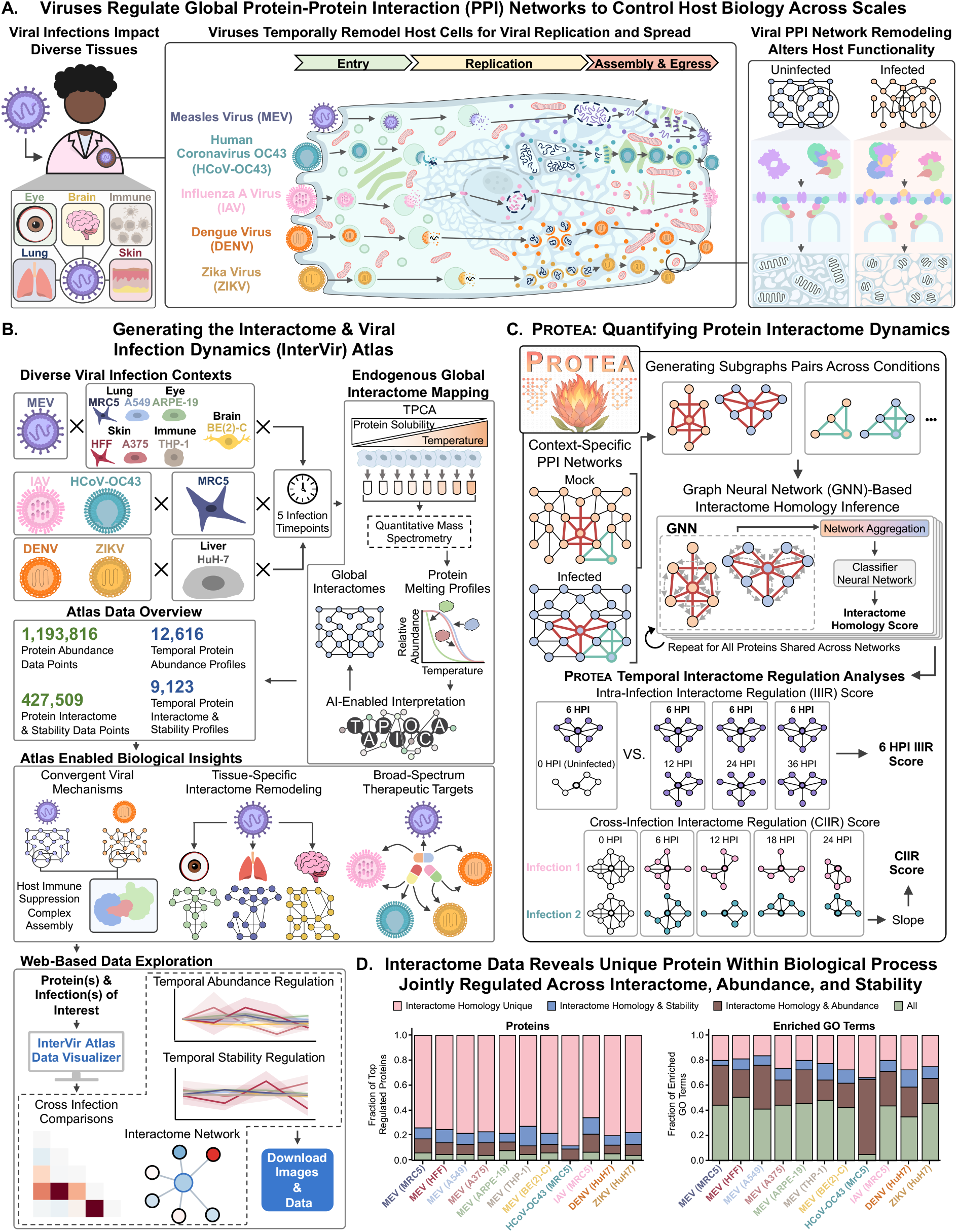
Generating the InterVir interactome atlas for RNA viral infections and developing PROTEA, a graph neural network-based AI framework for analysis of temporally resolved protein-protein interaction (PPI) networks. **A**, Viral infections regulate host biology across scales. By regulating PPI networks, viruses temporally and physically remodel host cells to facilitate their replication, as depicted for five RNA viruses (IAV, HCoV-OC43, DENV, ZIKV, and MEV). This regulation can vary across tissues and infection types. **B,** Schematic depicting the generation of Interactome & Viral Infection Dynamics (InterVir) Atlas, its data, and enabled biological insights. The InterVir Atlas contains global endogenous interactomes the five RNA viruses mentioned in **A** across five timepoints representing their replication cycles. Infections are performed in lung fibroblasts, liver, skin, immune, eye, and brain cells, as illustrated for the different viruses. To enable accessible exploration of data within the InverVir atlas, we developed the InterVir Atlas Data Visualizer website (https://interviratlas.princeton.edu/), allowing users to generate downloadable plots and perform temporal and comparative analyses. **C,** Schematic representation of PROTEA inputs, processing, output, and enabled downstream analyses. **D,** Stacked box plots depicting the overlap in regulated proteins via their interactomes with proteins regulated by abundance or stability (left), and the overlap in enriched GO terms across infections (right).

Mapping temporally resolved PPI networks across diverse viral infections is critical for uncovering both virus-specific replication strategies and shared regulatory hubs, the latter of which can inform of targets for broad-spectrum antiviral therapies. Expanding the scope of these measurements to multiple cell types can offer an improved understanding of viral tissue tropism and predict the efficacy of proposed therapies. To gain such insights into virus-host interactome networks, methods that can facilitate the characterization of global, temporal interactomes for endogenous proteins are valuable. Indeed, previous interactome studies have advanced our understanding of the biology of diverse types of viral infections^15–22^, and these investigations frequently relied on the expression of individual tagged viral proteins and sometimes were performed at single-infection time points.

A method that has proven valuable for capturing global endogenous interactomes during viral infections is thermal proximity coaggregation (TPCA) mass spectrometry^23–31^. Utilizing the principle proteins aggregate when exposed to temperatures^32–34^ and that interacting proteins denature together, TPCA uses similarities of resulting melting curves for predicting interactions for thousands of endogenous proteins^25,27,28^. We previously reported an AI framework, Tapioca^23^, that integrates TPCA derived protein melting curves with protein physical properties, domains, and tissue-specific functional networks^35^ to capture the dynamics of context-specific protein interactomes. In addition to mapping interactomes, TPCA additionally provides data on protein abundance and thermal stability, making it a powerful tool for assessing protein regulation across multiple dimensions within a single experiment. While we and others have used TPCA to reveal infection biology^23,24,27,28,36^, comparative analysis of these data-rich networks across multiple conditions remains a major computational bottleneck. TPCA datasets are noisy, and quantifying interactome shifts requires evaluating both changes in interactor identity and complex assembly/disassembly dynamics. Prioritizing which host proteins are most significantly altered across temporally resolved, multi-virus datasets is therefore non-trivial.

Here, we sought to enable interactome-based comparative virology studies and the identification of hubs for broad-spectrum antiviral therapies by generating an Interactome & Viral Infection Dynamics (InterVir) Atlas and developing a graph-based AI analytical framework, PROTEA. Our InterVir Atlas focuses on RNA viruses, leveraging TPCA to map temporal protein regulation across interactome, abundance, and thermal stability axes. We build interactome repositories for measles virus (MeV), influenza A virus (IAV), dengue virus (DENV), Zika virus (ZIKV), and human coronavirus OC43 (HCoV-OC43) infections across five infection timepoints and in relevant cell types for each virus. Collectively, these pathogens infect hundreds of millions of individuals annually^37–42^, yet approved antivirals are not yet available for MeV, DENV, ZIKV, and neither is there knowledge about potential broad-spectrum antivirals for treating a combination of these viruses. Further interrogating how host tissue/cell type impacts viral infection dynamics, the InterVir Atlas additionally contains temporally resolved data during MeV infection across seven cell types representing five tissues (i.e., skin, lung, immune, eye, and brain). To enable users to explore the InterVir Atlas, we developed a website (https://interviratlas.princeton.edu/) to allow data visualization for proteins of interests, comparison of infections, and downloading of figures and data for customized analyses. Spanning diverse genome organizations, tropisms, and pathogenesis profiles, the InterVir Atlas represents the largest temporal global PPI network dataset of viral infections to date.

To facilitate comparative virology investigations, in this study we introduce the concept of interactome homology. We developed PROTEA, a graph-based AI framework that quantifies proteome-wide interactome changes and discovers the similar protein interactome features across biological contexts. By analyzing temporally resolved interaction networks, PROTEA simplifies tracking protein interactome regulation and aids in the identification of meaningful interaction changes as the cellular environment is altered.

Our application of PROTEA for the systematic analysis of temporal interactomes within our InterVir Atlas facilitated both biological insights and the discovery of compounds that can be repurposed as antivirals across multiple infections. In MeV infections, PROTEA identified a tissue-specific MeV replication trajectory in a neuronal cell type. We found altered interactome regulation of the host signaling protein SHC1^43^, which we link to dysregulation of MeV gene expression. In DENV infection, we identified unique interactome regulation of alpha and gamma secretases and the thermal destabilization of selected cleavage targets. This DENV-specific regulation provides additional insights into molecular features that may underlie the vascular leakage associated with severe DENV infections^44^. Comparing all five infection types, we also uncovered hidden relationships between the studied viruses. We discovered that, at the interactome regulation level, MeV infection is most similar to flaviviruses, and particularly DENV, while MeV is most similar to IAV and HCoV-OC43 at the level of protein abundance regulation. Finally, following up on our PROTEA-facilitated discovery of convergent regulatory hubs across infections, we identified and experimentally validated ten small molecules that demonstrate efficacy in inhibiting multiple types of viral infections.

## RESULTS

### Constructing a Interactome & Viral Infection Dynamics (InterVir) Atlas

To identify conserved interactome remodeling across viral families and differential protein regulation across tissues, we coupled TPCA with Tapioca^23^ data processing to generate the largest to date temporal atlas of proteome-wide PPI networks during viral infections (Figure 1C). Our InterVir Atlas encompasses both inter-viral infections and inter-cell type comparisons. The atlas includes the global endogenous interactomes for five major RNA viral pathogens (IAV, HCoV-OC43, DENV, ZIKV, and MEV) across timepoints that represent their replication cycles. Comparisons are performed in relevant cell models (HCoV-OC43 and IAV in lung fibroblasts, and DENV and ZIKV in liver cells) at 0, 6, 12, 18, and 24 hours post-infection [HPI]. To expand consideration of cell type differences, MeV was selected as a model for broad tissue tropism. Hence, MeV infection was performed across seven cell lines representing lung (MRC5, A549), skin (HFF, A375), immune (THP-1), eye (ARPE-19), and brain (BE(2)-C), with each infection being monitored at five timepoints (0, 6, 12, 24, and 36 HPI). As MeV routinely infects lung, skin, immune, and eye tissues in vivo, but only rarely targets the brain^45–48^, this atlas enables the investigation of interactome regulation across both common and uncommon replication environments. Given that TPCA simultaneously captures protein abundance, thermal stability, and interactomes, the resulting atlas is exceptionally data-rich, comprising 55 global interactomes, alongside 12,616 protein abundance profiles and 9,123 protein stability and individual interactome profiles (Figure 1B; Supplementary Figures S1-S2).

To facilitate the exploration of this generated resource, we developed a data visualization website (https://interviratlas.princeton.edu/). This website allows researchers to specify both proteins and infections of interest to create plots that depict how these proteins are temporally regulated across protein interactome, abundance, and thermal stability dimensions. The plots generated and their data can be downloaded, enabling further custom downstream analysis. An overview of the website is provided (Supplementary Figure S3).

### PROTEA: An AI framework for the comparative analysis of context-specific PPI networks

Experimentally derived, proteome-wide PPI networks are information-rich resources for understanding protein function across cellular states^23–31^. However, the interpretation and comparison of these large interactomes are hindered by their scale, inherent noise, and the difficulty of interpreting how local interactome changes reflect meaningful functional shifts. These challenges amplify when moving beyond pairwise comparisons to analyzing temporal datasets across conditions, such as different viral infections.

To overcome these limitations, here we introduce PROTEA, a graph neural network (GNN)-based AI framework for the efficient, interpretable analysis of context-specific PPI networks (Figure 1C). Comparing networks on a per-protein basis, PROTEA quantifies interactome homology, a measure of protein interactome similarity, enabling the ranking of proteins by degree of interactome alterations between a pair of biological conditions. Representing proteins as a graph containing its local interactome and using with protein language model^49^ derived node embeddings to encode protein identify, PROTEA uses a graph neural network (GNN) to distinguish between conserved and divergent protein interactomes. The resulting PROTEA output, an interactome homology score, can be used to rank proteins by the extent of their interactome alteration, prioritizing candidates for downstream investigation. Trained on PPI networks from diverse biological conditions, including primary and cancer cells, viral infections, and different TPCA methodological variants^23–26,50,51^, PROTEA achieves a 5-fold cross-validation AUROC of 0.81 (Supplementary Figure S4A). The graph-based architecture facilitates interpretability, as proteins can be interrogated at the subgraph level to pinpoint the specific alterations, such as complex assembly/disassembly or interactor gain/loss, driving their score. As demonstrated below, PROTEA facilitates multi-scale analyses spanning individual proteins, complexes, and organelles. It readily identifies convergently regulated targets across diverse infections for therapeutic prioritization and uncovers regulatory mechanisms not captured by standard protein abundance or stability profiling alone.

Processing interactomes from our InterVir Atlas with PROTEA, we derived two temporal metrics (Figure 1C). *Intra-infection interactome regulation* (IIIR) quantifies the magnitude and consistency of interactome changes throughout a single infection time course. *Cross-infection interactome regulation (CIIR)* measures interactome convergence or divergence between pairs of infections. Specifically, a high IIIR score denotes a protein whose interactome shifts significantly from mock to infected states but remains largely stable across subsequent infection timepoints, suggesting its transition into a defined infection-associated state. A high CIIR score indicates that a protein’s interactome becomes increasingly similar between two distinct viral infections as they progress, highlighting potentially conserved functional roles.

To determine whether interactome regulation captures information orthogonal to protein abundance and stability, we assessed cross-modality correlations and found them to be negligible (Supplementary Figures S4B). Furthermore, 70-80% of the top interactome-regulated proteins were not flagged as highly regulated by abundance or stability metrics (Figure 1E). However, gene ontology (GO) enrichment analysis of these top proteins revealed that only 20-30% of enriched biological processes were unique to the interactome modality. Together, this indicates that while abundance, stability, and interactome shifts converge on the same core biological pathways, interactome profiling implicates distinct subsets of proteins, yielding orthogonal biological insights.

### MeV Infection Drives Cell Type-Specific Remodeling of Global PPI Networks

MeV has reemerged as a public health threat in the United States^37,52^, and, while effective vaccines exist^53^, there are no approved antivirals. A defining hallmark of MeV infection is systemic dissemination, which profoundly suppresses host immune function and drives high clinical severity relative to several other common respiratory viruses^45^. Given its broad tissue tropism, spanning lung, skin, immune, eye, and brain tissues, we investigated how consistently the host interactome is rewired across diverse cellular environments and whether localized differences impact viral outcomes. We generated and analyzed temporally resolved MeV interactomes across the seven corresponding cell types in our atlas. Using PROTEA, 105 pairwise comparisons (21 cell type pairs × 5 timepoints) allowed distilling large-scale network dynamics into interpretable, protein-level insights. Using IIIR and CIIR scores across cell types, we performed PCA, followed by Leiden clustering, and generated a UMAP visualization across MeV infections (Figure 2A; Methods). Cluster analysis revealed that only ∼20% of proteins displayed convergent regulation, with the vast majority exhibiting highly cell-line-specific interactome responses (Figure 2B). GO term enrichment of this convergent core highlighted well-established MeV biological processes, including mitochondrial energetics and morphology^54,55^, cell adhesion^56^, and viral genome replication^57^ (Figure 2C). Comparing the convergence between individual pairs of cell types revealed consistent regulation of actin filament polymerization and cell projection assembly across most cells (Supplementary Figure S5). Complementary analyses of cell-line-specific proteins highlighted the specific regulation of immune components, such as interferon-gamma and interleukin-8 production. Notably, in the BE(2)-C neuronal cell line, we observed unique interactome regulation of JUN and MAPK activity, the ERK1/2 signaling cascade, and innate immune signal transduction, processes known to be critical for MeV replication^58–60^. Therefore, we asked what specific proteins drove this distinct interactome regulation, finding that SHC1 was a core component for the MAPK enrichment.

**Figure 2.**
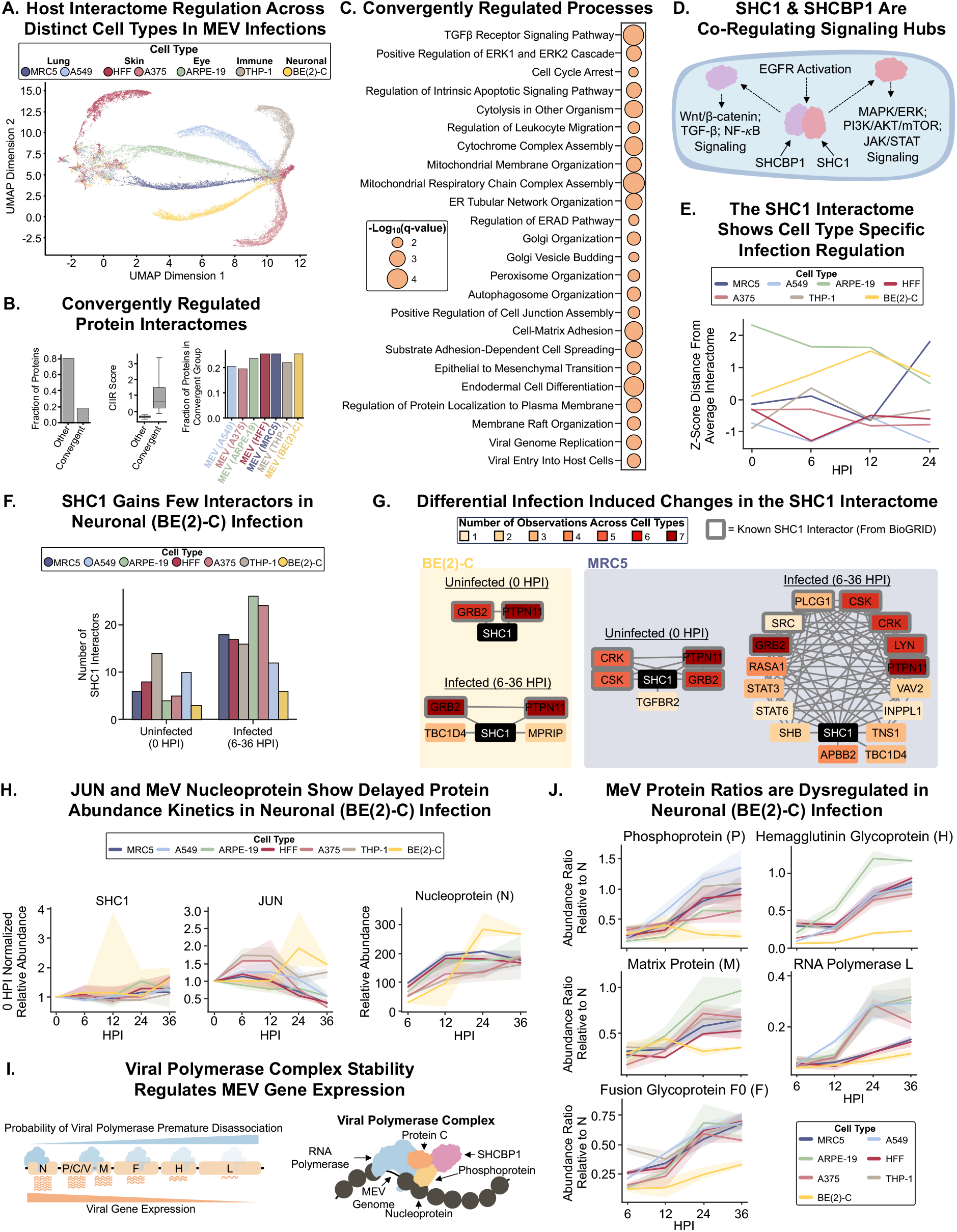
Cell type dependent MeV infection trajectories via the differential interactome regulation of SHC1 and the viral polymerase complex. **A**, UMAP of protein interactome regulation during MeV infection across seven cell types. In the plot, each dot represents a single protein in a specific, color denoted, cell type. **B,** Using Leiden clustering in PCA space, the bar chart depicts the fraction of proteins present in convergent or cell type specific clusters (*left*). Box plot showing the distribution of CIIR scores for proteins in convergent versus cell type specific clusters (*middle*). The median value (line) and the ±1.5 interquartile range (whiskers) are shown. Bar chart depicting the fraction of proteins from each cell type within the convergent clusters (*right*). **C,** GO term enrichment, performed using HumanBase^130,138^, of proteins whose interactomes display high cross-infection regulation across most of the examined cell types. **D,** Schematic depicting the roles and co-regulation of SHC1 and SHCBP1 in signaling pathways downstream of EGFR activation. **E,** PCA was performed on the IIIR and CIIR values for SHC1 across infections. For each timepoint, the mean PCA value was calculated and the distance from this mean point was computed for all infections. The Z-score normalized distances are shown in the line plot, illustrating the distance from the “average” SHC1 interactome per timepoint across infections. **F,** Bar plot showing the number of confident (see methods) SHC1 interactors in uninfected and infected states per cell type. **G,** SHC1 interactomes during uninfected and measles virus infected states for two of the tested cell types (see remaining cell types in Supplemental Figure 4B). In the interactomes, nodes are colored by the number of cell types that share the given interactor (counted separately between uninfected and infected interactomes), except for the SHC1 node, which is colored black. **H,** Temporal relative protein abundances, normalized to 0 HPI (hours post infection), of SHC1, JUN, and the MeV nucleoprotein during MeV infection across all seven cell types. The solid line represents the median value, and the shaded region represents the 95% confidence interval. **I,** Schematic showing the gene expression pattern of MeV genes regulated by the viral polymerase complex disassociation from the viral genome. **J,** Temporal abundances of select viral proteins per cell type normalized to respective viral nucleoprotein abundances. The solid line represents the median value, and the shaded region represents the 95% confidence interval.

While not previously studied in MeV infection, SHC1 is an important adaptor protein and hub for signaling across cellular proliferation, survival, immune response, and cytoskeletal remodeling^43,61^. Central for these functions is the interaction of SHC1 with its binding partner, SHCBP1, which sequesters both proteins to the cytoplasm. If EGFR signaling is activated, this complex dissociates and the two proteins are freed to function within signaling pathways, including MAPK/ERK, TGF-β, Wnt/β-catenin, PI3K/Akt, and JAK/STAT signaling^43,61–63^ (Figure 2D). Given this broad functionality and flagging of SHC1 as uniquely regulated in MeV infection in the tested neuroblastoma cells, we further characterized the temporal SHC1 interactomes across MeV infections. Using SHC1 IIIR and CIIR scores in PCA space, we computed the distance between cell type-specific SHC1 interactomes across infection timepoints (Figure 2E). The eye SHC1 interactome was unique in both uninfected and infected states compared to other cell types. In contrast, the neuronal SHC1 interactome displayed infection-specific features. While being similar for most cell types prior to MeV infection, the neuronal SHC1 interactome diverged from other cell types during infection. These findings suggest neuronal- and eye-specific regulation of SHC1 in MeV infection.

Further assessing the SHC1 interactomes, we observed an expansion in the number of interactions upon infection in most cell types, with the exception of the neuroblastoma cell type (Figure 2F; Supplementary Figure S6A). Across most cell types, the retained SHC1 interactors included known partners, such as PTPN11 and GRB2^43^ (Figure 2G, Supplementary Figure S6B). Upon infection, SHC1 broadly recruited known and novel partners linked to MAPK/ERK signaling, PI3K/AKT/mTOR pathways and cytoskeletal organization (RASA1, LYN, TNS1)^64–66^ in a cell type specific manner. While the infected neuronal SHC1 gained a cytoskeletal organization related interactor, MPRIP^67^, it gained no additional interactors relating to signaling pathways. Given our observation of an expanded SHC1 interactome in most cells, we considered the possible contribution of altered protein abundance (Figure 2H). While SHC1 displayed similar abundance trends across cell types, JUN, its downstream transcription factor involved in immune responses^43,68^, had cell type-specific temporal regulation. In most cell types, JUN abundance peaked early (6 to 12 HPI), then declined late in infection (24 to 36 HPI). In neuronal cells, this peak was delayed to 24 HPI, while in eye cells, abundance declined from the start of infection. These distinct temporal patterns may be linked to altered infection kinetics or a broader dysregulation of infection in neuronal cells. As a proxy of infection kinetics, we analyzed the temporal abundance of the MeV nucleoprotein (Figure 2H), which controls the shift of the viral polymerase complex from transcription to genome replication^69^. Nucleoprotein abundance increased before plateauing after 12 HPI in most cells, but this trend was delayed in neuronal cells. Together with our interactome results, these observations suggest a possible connection between SHC1 regulation and MeV infection kinetics.

MeV gene expression occurs along a gradient, driven by the unstable association of the viral polymerase complex with the viral genome^48,57^. This results in higher relative protein abundances for genes located near the 3’ end relative to genes near the 5’ end (e.g., high nucleoprotein N, low RNA polymerase L) (Figure 2H). The SHC1 partner, SHCBP1, has been shown to interact with the viral polymerase complex, stabilizing its association with the viral genome and ensuring viral proteins are produced in the correct ratios^70^. Given the limited alterations in SHC1 interactions and the delayed JUN and MeV nucleoprotein expression in neuronal cells, we hypothesized that the SHC1-SHCBP1 dissociation may be delayed or otherwise damped in MeV infection in this cell type. If this is the case, then we would expect viral proteins to be expressed in incorrect ratios, particularly for proteins encoded by genes closer to the 5’ end of the viral genome. Indeed, our results showed dysregulated temporal ratios of viral proteins compared to nucleoprotein (Figure 2I; Supplementary Figure S6C).

We next explored other possible explanations for these differential viral protein ratios in neuronal cells, considering both other host and viral factors. Monitoring another host regulator of MeV gene expression, HSP72 (HSPA2)^57^, we observed convergent HSP72 interactome regulation across most cell types (Supplementary Figure 6D). Turning our attention to viral members of the polymerase complex, we found strong interactome conservation for the RNA polymerase L across all, and for photoprotein across most, cell types (Supplementary Figure 6E). The viral protein C, also involved in stabilizing viral polymerase complex and viral genome association, showed interactome regulation divergence across most cell types. However, the protein C interactomes were similar across neuronal, lung, and immune cell types, not reflecting the differential viral protein abundance regulation. These observations suggest that the SHC1 interactomes more closely align with the differential regulation of viral protein ratios than these other host and viral proteins. Additionally, these results highlight the value of our InterVir Atlas and PROTEA framework for disentangling how cell type differences impact viral infection outcomes.

### Identifying Convergent Cellular Remodeling Across Respiratory Viruses and Arboviruses

Beyond individual cell type variations, our InterVir Atlas enables the systematic detection of conserved interactome rewiring across distinct viral lineages. This atlas encompasses respiratory pathogens (MEV, HCoV-OC43, IAV) and arboviruses (DENV, ZIKV), representing an array of genetic architectures, positive-sense (HCoV-OC43, DENV, ZIKV), negative-sense (MeV), and segmented negative-sense (IAV) single-stranded RNA viruses ^1^. These pathogens also utilize distinct subcellular replication sites (nucleus for IAV; ER for HCoV-OC43, DENV, ZIKV; cytoplasm for MeV)^5,71–73^ and induce clinical outcomes spanning from predominantly mild (HCoV-OC43, ZIKV)^74,75^ to potentially life-threatening (MeV, IAV, DENV) diseases^45,76,77^. These divergent phenotypes are driven by cellular remodeling across multiple molecular modalities, and are captured in temporal alterations in protein abundances, stability, and interactome networks. Using PROTEA to analyze these diverse infections, we sought to uncover shared viral mechanisms of cellular remodeling. Performing PCA clustering and UMAP visualization across all five viruses, we establish that most proteins exhibited infection-specific interactome dynamics (Figures 3A-B). GO enrichment of the convergent protein subset highlighted pathways central to viral lifecycle requirements, physical organelle remodeling (mitochondrial/peroxisome fission, ER-to-Golgi transport), cell cycle arrest, and apoptosis (Figure 3C). Pairwise infection comparisons further enriched for energy production, oxidative stress, peptidase activity, membrane organization, cytoskeletal reorganization, and immune responses (Supplementary Figure S7).

**Figure 3.**
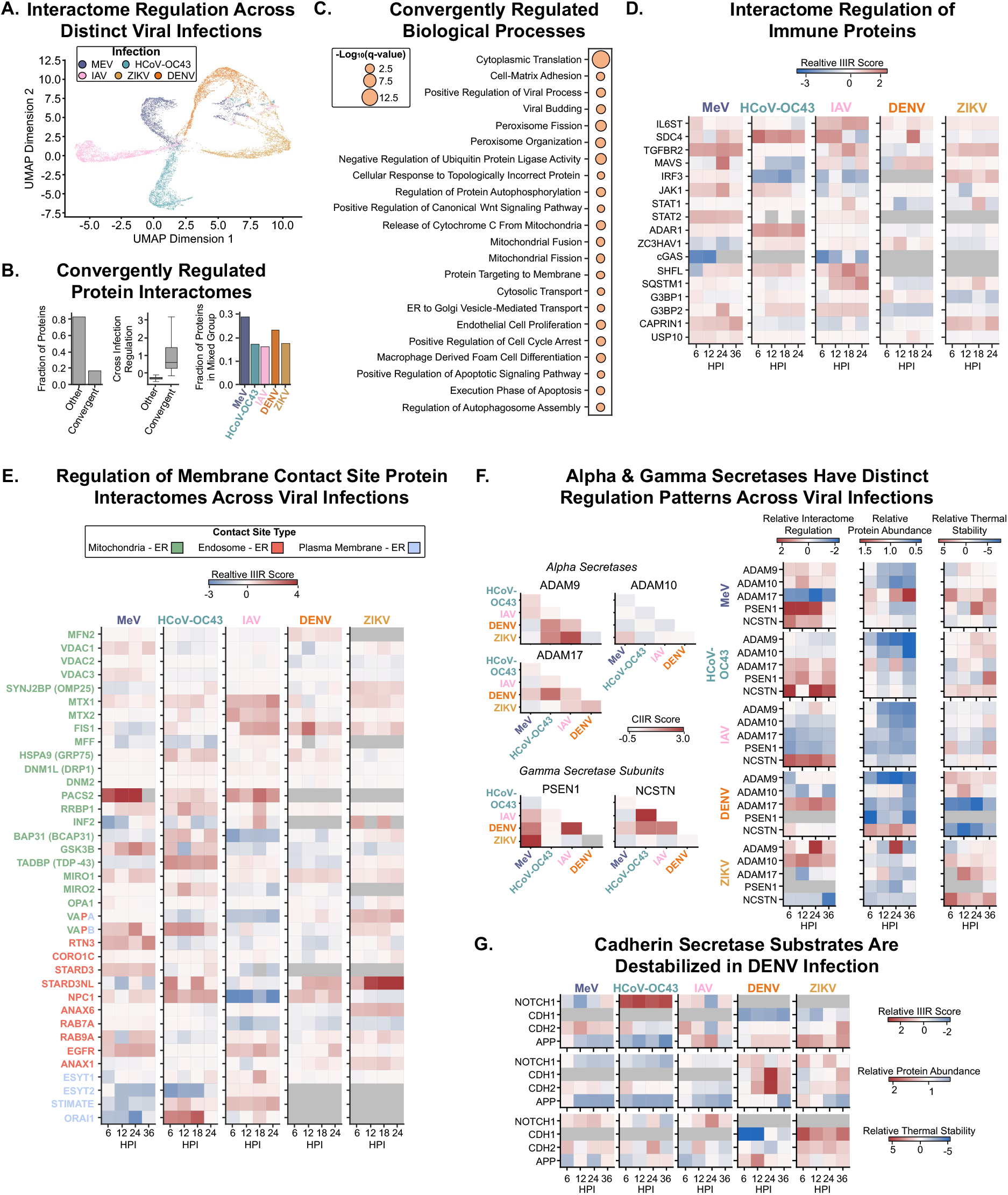
Respiratory and flaviviruses show distinct patterns of convergence and divergence across immune, membrane contact site, and secretase regulation. **A**, UMAP of protein interactome regulation during MeV, IAV, and HCoV-OC43 infections in fibroblasts, and DENV and ZIKV infections in liver cells. In the plot, each dot represents a single protein in a specific, color denoted, infection. **B,** Using Leiden clustering in PCA space, the bar chart depicts the fraction of proteins present in convergent or infection specific clusters (*left*). Box plot showing the distribution of CIIR scores for proteins in convergent versus infection specific clusters (*middle*). The median value (*line*) and the ±1.5 interquartile range (whiskers) are shown. Bar chart depicting the fraction of proteins from each infection within the convergent clusters (right). **C,** GO term enrichment, performed using HumanBase^130,138^, of proteins whose interactomes display high cross-infection regulation across most of the examined infections. **D,** Heatmaps depicting the Z-score normalized IIIR scores of immune related proteins across infections and timepoints. **E,** Heatmaps depicting the Z-score normalized IIIR scores of membrane contact site proteins across infections and timepoints. **F,** Heatmaps depicting the CIIR scores between infection pairs, Z-score normalized IIIR scores, 0 HPI normalized protein abundance, and Z-score normalized thermal stability for alpha and gamma secretase proteins across infections and timepoints. **G,** Heatmaps showing the Z-score normalized IIIR scores, 0 HPI normalized protein abundance, and Z-score normalized thermal stability for protein substrates of alpha and gamma secretases across infections and timepoints.

To delve more deeply into our integration of the interactome, abundance, and stability measurements, we started by evaluating the regulation of immune-related proteins (Figure 3D; Supplementary Figures S8A-C). When considering protein abundance changes, as positive controls, we observed the previously reported increase in STAT1 during IAV infection^78^, the degradation of STAT2 in ZIKV infection^79^, as well as the increase of SHFL abundance in ZIKV, DENV^80^, and IAV infections^78^. Other immune factors exhibited differential regulation at the abundance, interactome, and stability levels. For example, the cytokine receptor IL6ST^81^ demonstrated convergent interactome regulation across most infections (except HCoV-OC43). However, its abundance trajectories were split, decreasing in MeV, HCoV-OC43, and IAV, while increasing in DENV and ZIKV infections. Another example comes from MAVS, a hub for interferon signaling^82^, which these viruses are known to regulate directly^83^ or via protein interactions upstream and downstream of MAVS signaling^79,84,85^. MAVS maintained its abundance across all infections but exhibited increased thermal stability and convergent interactome regulation during MeV, ZIKV, and DENV infections.

Given that physical organelle remodeling featured prominently in our convergent GO analysis, we next examined membrane contact site (MCS) proteins, which mediate organelle morphology and inter-organelle communication (Figure 3E; Supplementary Figures S9A-C). While IAV infection moderately decreased the abundance of most MCSs, aligning with previous reports^86^, PROTEA uniquely identified strong interactome regulation of MTX1, MTX2, and FIS1, which regulate mitochondrial fission^87^, a known phenotype of IAV infection^88^. For MeV, we observed profound interactome dysregulation at plasma membrane-ER contacts. Specifically, the lipid transporter ESYT2^89^ increased in abundance, while the calcium channel ORAI1^90^ increased in thermal stability. These changes likely facilitate syncytia formation late in MeV infection, where localized calcium influx (via stabilized ORAI1) and lipid transport to expanding membranes (via upregulated ESYT2) are required for massive cell-to-cell fusion^91,92^. Broadening this to convergent organelle remodeling, the late endosome-ER tethering protein STARD3NL^93^ exhibited highly conserved interactome regulation across most infections, despite possessing divergent abundance and stability profiles across the atlas.

Our GO term analysis also enriched for “positive regulation of peptidase activity,” drawing our attention to alpha and gamma secretases, which are known mediators of viral entry and pathogenesis (Figure 3F). Among alpha secretases^94^, ADAM9 and ADAM17 displayed broad interactome convergence. However, we noted virus-specific deviations, with ADAM9 and ADAM10 being strongly regulated in ZIKV and MeV infections, whereas ADAM17 dominated in DENV infection. Therefore, closely related flaviviruses (DENV and ZIKV) may modulate secretase in a distinct manner. Gamma-secretase subunits PSEN1 (catalytic) and NCSTN (receptor)^95^ similarly showed complex, virus-specific interactome and abundance regulation, predominantly marked by global downregulation across most infections, except during DENV infection, where NCSTN abundance increased.

To understand the downstream consequences of the secretase rewiring, we examined their known cleavage targets (Figure 3G). NOTCH1, cleaved by ADAM10, ADAM17, and gamma-secretase^96,97^, exhibited a distinct interactome shift in HCoV-OC43 infection, directly mirroring the unique ADAM17 regulation observed for this infection. Furthermore, the cell adhesion proteins CDH1 and CDH2 (cleaved by ADAM10 and gamma-secretase)^98,99^ increased in abundance but decreased in thermal stability during DENV infection. This thermal destabilization may indicate a failure of these proteins to integrate into functional adhesion complexes, likely due to altered secretase regulation. In contrast, ZIKV infection increased the stability of these cadherins. To further map alpha secretase divergence, we analyzed the interactomes of these enzymes across infections. During MeV, HCoV-OC43, and IAV infections, these secretases displayed a convergent core interactome. This included interaction with additional alpha secretases and engagement with secretory and folding machinery, such as CANX, CPD, LMAN1^100–102^. In contrast, the flaviviruses were more distinct, with ZIKV having a small, and DENV a more extensive, alpha secretase interactome.

Both DENV and ZIKV disrupt intercellular junctions via circulating, secreted viral NS1 proteins^103–105^; however, while ZIKV induces subtle junctional alterations to cross the blood-brain barrier^106^, DENV frequently causes severe, life-threatening systemic vascular leakage^44,107^. Our data provides a bipartite mechanistic explanation for this clinical phenotype. First, the DENV alpha secretase interactome (Supplementary Figure S10) directly targets cytoskeletal and junctional complexes (e.g., ENAH, DSC2)^108,109^ alongside extracellular matrix modulators (e.g., EDIL3)^110^. Acting in concert with the observed CDH1/CDH2 destabilization, this physically dismantles the endothelial barrier while potentially preventing leukocyte adherence. Second, this expanded network uniquely sequesters critical immune regulators. The incorporation of danger-associated molecular patterns (DAMPs, like HMGB2)^111^ and immune regulators (like RIOK3)^112^ likely contributes to cytokine storms, while simultaneously blunting early interferon responses. This immunomodulation is compounded by the DENV-specific regulation of ADAM17 (Figure 3F), an important factor in the secretion of TNF-α and other key inflammatory cytokines^113^. Ultimately, by coupling hyper-inflammatory signaling and localized immune evasion with the direct structural degradation of adhesion complexes, this dual mechanism can explain why DENV triggers profound vascular leakage unlike the other viruses studied here^44^ .

Together, these findings demonstrate the power of integrating interactome, abundance, and thermal stability dynamics. By tracking multi-modal regulation across functionally related protein networks, the InterVir Atlas not only highlights mechanistic divergence between even closely related pathogens but also identifies the shared host dependencies critical for pan-viral therapeutic intervention.

### Repurposing drugs and small molecules as antivirals for multiple viral infections

Many human viral pathogens still lack approved antiviral therapies, which is also the case for some of the viruses explored in this study (i.e., MeV, HCoV-OC43, DENV, and ZIKV). Additionally, given the frequent difficulty in identifying the type of infectious virus affecting an individual, there is considerable interest in developing broad-spectrum antivirals^114^. We hypothesized that PROTEA-derived infection interactome scores can help address this challenge (Figure 4A). We sought to map our PROTEA scores onto a library of drugs and small molecules known to target protein within our identified regulatory hubs. Hence, this analysis can help prioritize broad-acting antiviral candidates even for compounds not previously used for treating viral infections.

**Figure 4.**
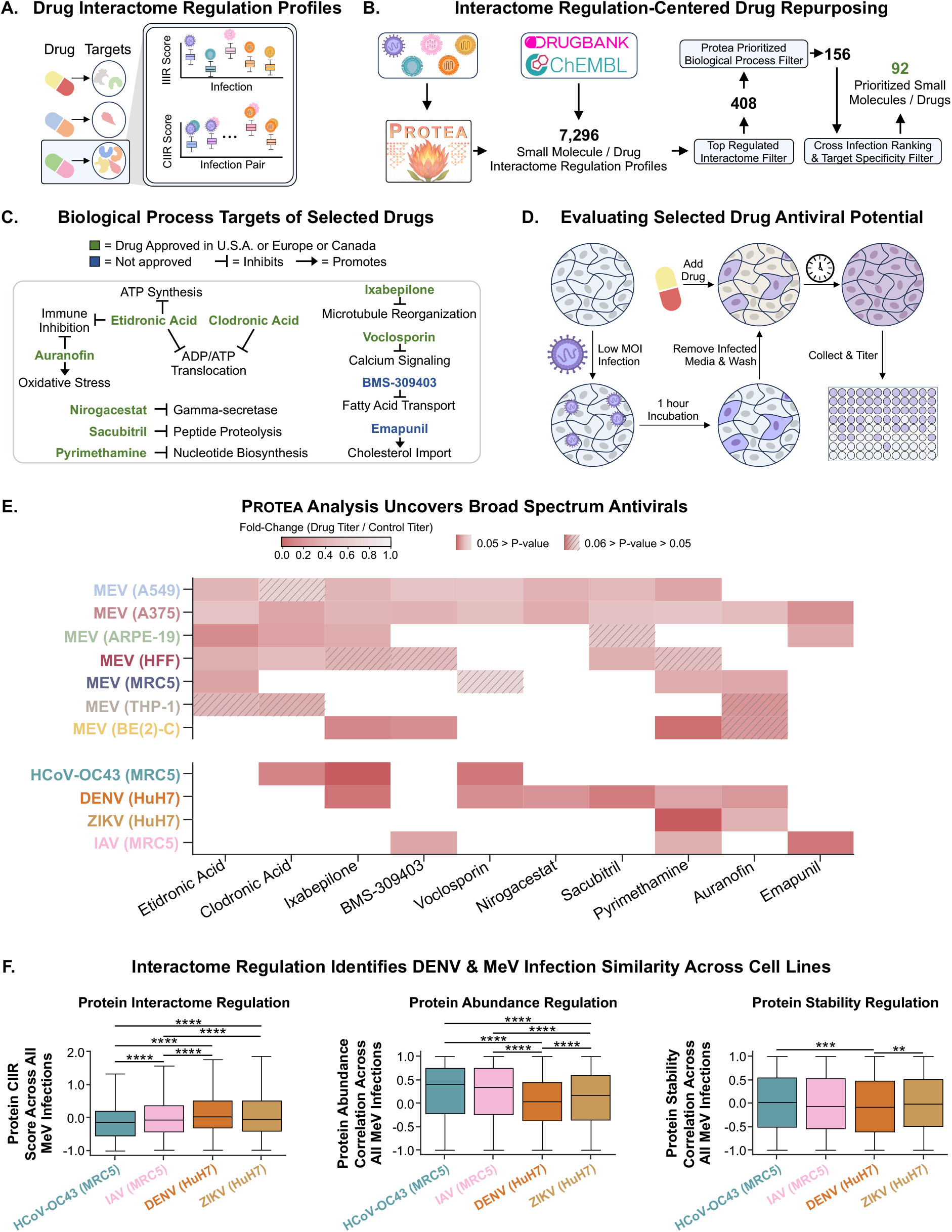
Leveraging PROTEA predictions to repurpose drugs and small molecules as antivirals effective across multiple infections. **A**, Schematic depicting the generation of drug interactome profiles. For each drug, the CIIR and IIIR scores for all known protein targets of the drug are aggregated to generate the drug interactome profile. **B,** Integration of PROTEA predictions with ChEMBL^115^ and DrugBank^116^ databases to generate 7,296 drug interactome regulation profiles. These were then filtered down to a list of 92 drugs and small molecules prioritized for repurposing as antivirals (see methods for more details). **C**, Overview of the ten drugs (manually selected from the list of 92) used in follow-up experiments. **D,** Schematic depiction of the experimental setup for evaluating the effect of the selected drugs on viral infections. **E,** Heatmap showing the viral titer fold-change in drug treatment versus control. For clarity, only conditions in which a p-value less than or equal to 0.06 was observed are shown; Supplementary Figure S11 contains all data. P-values were calculated using a one-sided Student’s t-test and adjusted for multiple comparisons using the Benjamini-Hochberg procedure. All data shown represents the average of three replicates. See Supplementary Figure S11 for p-values and individual plotting of data points. **F**, Box plots showing the CIIR score (*top*), protein abundance correlation (*middle*), or protein stability correlation (*bottom*) with MeV infections of top regulated proteins by IIIR score, protein abundance, or thermal stability, respectively, in HCoV-OC43, IAV, DENV, and ZIKV infections. The line within the box represents the median value and the whiskers represent the ±1.5 interquartile range. P-values were calculated using a two-sided student’s T-test and were adjusted for multiple comparisons using the Benjamini-Hochberg procedure. Only p-values ≤ 0.5 are shown and are represented as * ≤ 0.05, ** ≤ 0.01, *** ≤ 0.001, and **** ≤ 0.0001.

Integrating PROTEA predictions with ChEMBL^115^ and DrugBank^116^, we generated 7,296 drug and small-molecule interactome regulation profiles. We systematically filtered these based on interactome regulation magnitude, relevant GO annotations, and cross-infection hit frequency, yielding 92 prioritized candidates (Figure 4B; Methods). From this pool, we selected 10 compounds for experimental validation based on their mechanistic plausibility, narrow biological process scope, and governmental drug approval status (Figure 4C). All 10 compounds demonstrated statistically significant antiviral activity against at least two infections. Viral titers were assessed following multicycle infections using low-multiplicity of infection (MOI) and measured using the 50% Tissue Culture Infectious Dose (TCID50) assay following 90% or greater cytopathic effect (CPE) in controls (Figures 4D-E; Supplementary Figure S11). None of the tested compounds exhibited significant cytotoxicity across the evaluated cell types (Supplementary Figure S12).

Amongst the compounds tested, those widely effective spanned diverse pharmacological classes, including bisphosphonates (etidronic acid, clodronic acid)^117,118^, chemotherapeutics (ixabepilone)^119^, antimalarials (pyrimethamine)^120^, and antirheumatics (auranofin)^121,122^. Notably, pyrimethamine and auranofin were previously identified in a large, untargeted screen of FDA-approved drugs in ZIKV infection^123^. While not tested for other viral infections, this overlap provides orthogonal validation of our PROTEA-driven computational prioritization. These results establish that temporally resolved interactome dynamics can successfully guide drug repurposing.

Our pipeline intentionally overrepresented MeV datasets, yielding seven drugs effective across multiple MeV infections in different cell types. Across these seven compounds, four were also effective for multiple other viral families. This cross-viral trend was even more pronounced for the six drugs effective against DENV. This was unexpected, given that MeV (a paramyxovirus) and DENV (a flavivirus) possess fundamentally distinct genome architectures, replication sites, and clinical pathologies^45,73,76^. We hypothesized that our temporal interactome analysis captured a hidden conserved regulatory axis linking these seemingly disparate viruses.

To test our hypothesis of a conserved infection programming, we analyzed proteins exhibiting the strongest interactome regulation (IIIR scores) across the non-MeV infections and compared their convergence to the seven MeV time courses (Figure 4F). Remarkably, DENV exhibited the highest cross-infection interactome regulation (CIIR) score distribution, significantly exceeding those of the respiratory viruses HCoV-OC43 and IAV. ZIKV similarly displayed a statistically significantly higher CIIR distribution than the respiratory viruses. These findings reveal that the MeV interactome regulation more closely mirrors those of flaviviruses than its respiratory counterparts. Protein abundance data illustrated the opposite relationship, with abundance changes in HCoV-OC43 and IAV correlating significantly more with MeV than with DENV or ZIKV. Further highlighting that the convergence can primarily be seen at the interactome level, the thermal stability profiles showed no clear alignment with either modality. This interactome-specific linkage persisted when resolving the analyses at the individual cell type MeV infection level (Supplementary Figure S13). Altogether, our results demonstrate that PROTEA-driven analysis of the InterVir Atlas successfully prioritized 10 actionable compounds for multi-viral drug repurposing, while uncovering a cryptic regulatory convergence between MeV and DENV. These findings highlight that global interactome dynamics provide a profound, orthogonal dimension to comparative virology, revealing targetable, functional relationships between pathogens that remain hidden to standard abundance and stability profiling.

## DISCUSSION

The dynamic remodeling of host protein-protein interaction (PPI) networks plays important roles across viral infections^11,12,14–18,23,24,27,28^. Mapping these events is critical for understanding the biology of infections, conserved vulnerabilities across distinct pathogens, and targets for broad-spectrum antiviral interventions. To this end, we leveraged TPCA to generate the largest-to-date, temporally resolved atlas of global host-virus PPI networks, monitoring five distinct viral infections across eight cell types. To promote the accessible and wide-spread dissemination of these data, we developed a data visualization website (https://interviratlas.princeton.edu/) for the exploration of the InterVir Atlas.

To systematically interpret our InterVir Atlas, we developed PROTEA, an AI framework utilizing graph-based deep learning to enable proteome-wide comparative analysis of context-specific PPI networks. By calculating interactome homology scores, PROTEA quantifies shifts in protein neighborhoods across biological contexts, systematically ranking the most profoundly rewired host proteins. Applied to our viral atlas, PROTEA successfully generated mechanistic hypotheses, explained how tissue-specific interactome regulation may drive organ-level pathogenesis, and accurately prioritized compounds for antiviral repurposing.

In MeV infection, PROTEA identified SHC1 as a regulator of viral protein ratios. Specifically, we found divergence in the SHC1 interactome regulation in neuronal and retinal cell types compared to the other cell types studied. Within the MeV neuronal infection, SHC1 failed to gain interactors with known signaling partners as was observed in all other cell types. We show that this regulation is not at the protein abundance level but rather is specific to protein interactions. Indeed, supporting this differential regulation, we identified delayed expression kinetics of JUN, a protein downstream of SHC1 signaling. Furthermore, linking this differential regulation of host factors to MeV replication outcomes, we identified delayed MeV nucleoprotein expression kinetics and altered MeV protein ratios specifically in the neuronal cell type. By sequestering SHCBP1, SHC1 can influence the stability of the viral polymerase complex on the viral genome, with its differential interactome regulation in neuronal and retinal cells likely driving the stark tissue-specific differences in viral gene expression. This dysregulation was particularly pronounced in the tested neuroblastoma cell type, having likely clinical implications. In the brain, MeV can cause subacute sclerosing panencephalitis (SSPE), a rare, almost always fatal condition that emerges up to a decade post-infection^124^. SSPE is characterized by a slow, defective viral spread, driven by genome mutations acquired during viral persistence^46,47,125^. These mutations cause aberrant viral protein ratios, similar to those we observed in our data. The fact that a single replication cycle of a non-neuron-adapted MeV strain induces this dysregulation suggests that the baseline neuronal host environment inherently predisposes the virus to this trajectory, potentially priming MeV to acquire the mutations necessary for chronic brain infection.

In the mammalian brain, the neuronal SHC1 expression is known to be age-dependent, with young neurons expressing high levels and adult neurons expressing little to none SHC1^126^. Clinically, the vast majority of SSPE cases occur in patients who contracted MeV before two years of age^127^. It is therefore plausible that the combination of elevated SHC1 expression and a neuron-specific SHC1 interactome leads to viral dysregulation, promoting persistent infection and subsequent mutagenesis characteristic of SSPE. Since neuroblastoma lines, like BE(2)-C, naturally express high levels of SHC1^128,129^, our results may at least partially recapitulate this SHC1 regulation. This effect of SHC1 is likely contributed to by multiple factors, including its interactome within a specific cellular type and context, given that the other cell types that we tested have even higher levels of SHC1 (Supplementary Figure S14). This underscores the immense value of cross-tissue interactome profiling and demonstrates the ability of PROTEA to decode rare, tissue-specific functions and resulting pathologies.

Broadening our PROTEA-driven analysis across the entire InterVir Atlas, we identified both unique regulatory patterns underlying specific mechanisms of infection and conserved hubs suitable for therapeutic targeting. For example, we identified a DENV-specific interactome and thermal stability shift for cadherins (CDH1/CDH2). The specific dysregulation of secretase-mediated cadherin cleavage we found offers further insights into how the virus systematically dismantles intercellular junctions, a phenotype notably absent in other infections evaluated in this study. This may provide additional insights into mechanisms underlying the known severe systemic vascular leakage induced by DENV infection^44,103,107^.

In contrast to these virus-specific mechanisms, we also made the unexpected observation that, at the interactome level, the flaviviruses DENV and ZIKV are similar to the paramyxovirus MeV. The respiratory viruses HCoV-OC43 and IAV shared greater similarity with MeV only at the level of protein abundance. Translating these convergent interactome insights into actionable therapeutic strategies, we demonstrate that PROTEA can successfully prioritize broad-spectrum antiviral candidates by mapping interactome regulation profiles onto drugs/small molecules. Our experimental validation of these predictions showed that all ten tested compounds are effective across multiple infection contexts. These results highlight that targeting conserved host-pathogen interactome dynamics, rather than viral proteins that can be highly mutated or only by focusing on changes in protein abundances, provides a highly efficient pipeline for broad-spectrum antiviral discovery.

While this study primarily highlights the mechanistic insights derived from interactome dynamics, our temporally resolved InterVir Atlas also contains over one million protein abundance and 400,000 thermal stability and interactome data points, providing an unprecedented resource for comparative virology. Furthermore, while applied here exclusively to viral infection, PROTEA is a fundamentally agnostic AI framework, readily adaptable to studying interactome regulation across other biological contexts. Ultimately, by illuminating the deep convergence of cellular remodeling across diverse infections, this computational framework and experimental atlas provide powerful tools for both fundamental biology and infectious disease preparedness.

## ACKNOWLEDGEMENTS

We thank all members of the Cristea and Troyanskaya laboratories for helpful discussions. We are grateful for funding from the NIH NIGMS (R01GM114141, I. M. C.), NIAID (AI174515, I. M. C.), Stand Up To Cancer Convergence (3.1416, I. M. C.), Paul Allen Foundation (I. M. C.), the CHDI Foundation (I. M. C.), the Gates Foundation (INV-081342) to O.G.T and I.M.C, the National Science Foundation Graduate Research Fellowship Program under Grant No. DGE-2039656 (awarded to T. J. R.), and a Moore Postdoctoral Fellowship (GBMF 13897) to T.J.R. Any opinions, findings, and conclusions or recommendations expressed in this material are those of the authors and do not necessarily reflect the views of the funding agencies. The funders had no role in study design, data collection and analysis, or the decision to publish or prepare the manuscript.

## AUTHOR CONTRIBUTIONS

T.J.R., O.G.T., and I.M.C designed the research. T.J.R. performed all experiments, method development, and computational analysis. T.J.R and A.T. developed the InterVir Atlas data visualization website. T.J.R wrote the initial manuscript, and T.J.R., O.G.T., and I.M.C edited and proofread the manuscript. O.G.T., and I.M.C supervised the project and obtained funding support.

## METHODS

### PROTEA Framework Details

#### Data preprocessing

The primary raw input into PROTEA are protein-protein interaction (PPI) network files that contain both the edges between nodes and associated edge weights. These network files contain a fully connected network. These networks are processed through several steps. First, to manage the number of possible downstream comparisons between networks, and to improve the confidence of predicted edges, PPI networks representing biological replicates of the same condition are combined by taking the median edge weight across replicates for a given edge for all edges. Edges only present in one replicate are preserved. Following network merging, multiple z-scores are computed from the edge weights. The first z-score is global, computed by using all edge weights present in the entirety of the network. The second z-score is local, calculated per protein by looking at only the edge weights associated with an edge containing the given protein of focus. The original edge weight, the global z-score, and the two local z-scores (one per protein participating in the edge) are used as edge embeddings. After calculating edge embeddings, edges are next pruned to transform the network from a fully connected network into a sparse network. First, edges whose edge weight values are below the 70^th^ percentile are pruned. The remaining edges are then filtered again pruning edges whose edge weight is below the 80^th^ percentile of the local edge weight distribution (local in that only edge weights from edges related to the protein are considered), unless this would leave a node with less then 20 edges, in which case this second level of filtering is skipped. Edges were also preserved through this second filtering step if the edge weight was in the 99^th^ percentile of all edge weights prior to 70th percentile filtering. These exceptions are included to ensure that all nodes retain some level of connectivity with the rest of the network and so that highly weighted interactions are not lost. Following this, a final layer of filtering is applied to prevent nodes from retaining an extreme number of edges which would be too large to process through the graph neural network portion of PROTEA. In this pruning step, all nodes are iterated through and edges are ranked by the degree of the connecting node (the node not currently in focus). The edge connected to the node of the highest degree is dropped sequentially until the node in focus has exactly 300 edges, the maximum allowed number of edges. The other raw input is averaged protein sequence embeddings from the ESM2 15 billion parameter model^49^. To acquire embeddings for proteins whose sequences were longer than the allowed sequence length for ESM2 we separated such proteins into chunks of 1,024 amino acids with 100 amino acids overlapping between adjacent chunks. Each of these chunks were then processed by ESM2. The average embedding across all amino acids was computed per chunk, and then a weighted average (weighted by the number of amino acids in per chunk) of all chunks average embeddings were then computed and used as the final embedding for use in PROTEA for the given protein. These processed embeddings are then used as node embeddings within the network.

#### Predicting interactome homology

After all preprocessing steps are completed, pairs of global PPI networks are generated for all possible nonredundant pairs of networks. For each protein, denoted the target protein, that is present in both networks within a pair, local PPI networks are generated. These local networks consist of the target protein, its interactors, and the edges between this set of interactors. Thus, the local networks denote the local interactome of a target protein and the state of that interactome. For a pair of local networks, representing the same target protein across two biological conditions, each network is individually processed through a graph autoencoder followed by pooling into a fixed-sized representational embedding. The representational embeddings from the pair of local networks are then concatenated and undergo a second round of pooling followed by fully connected linear layers for final classification. More specifically, the graph autoencoder consists of three blocks, themselves each containing three sub-blocks of PDNConv layer^131^ followed by leaky ReLU activation followed by a dropout layer used during training. Within the sub-blocks, the number of in and out channels remain constant. For the main blocks, the number of in and out channels are constant for the first two blocks, and the number of output channels for the third block is halved. The output of the first block is processed by a second block, with its features integrated via a residual skip connection. To facilitate deeper feature extraction while maintaining dimensionality, the subsequent stage employs a projected residual connection, wherein the input to the third block is transformed through a linear layer before being aggregated with the block’s output. Global pooling on the graph is performed using memory-based pooling^132^ and is followed by layer normalization after each pooling step. After the pooling steps, two linear layers are used for the final binary classification prediction. The PROTEA model was constructed using the PyTorch^134^ and PyTorch Geometric^136,137^ libraries.

#### Training, testing, and validation

The PROTEA framework was trained using a two-stage transfer learning approach. In the first stage, the graph autoencoder portion of the model was pretrained in an unsupervised manner to learn latent representations of the graph structure through predicting masked edges on local graphs. In the second stage of training, the graph autoencoder weights were frozen, and the pooling and linear layers were trained for binary classification using a binary cross entropy loss function. For classification training, global context specific PPIs generated from TPCA data were sourced from diverse biological contexts^23–26,50,51^ (using only data from control experiments from these datasets). To generate positive examples from these data, first the distance between and standard deviation of melting curves for the same across replicates for all datasets was computed. Then, per dataset, we took proteins whose curve replicates were in the bottom 20^th^ percentile for both distance and standard deviation as true positive examples. This was done with the expectation that proteins with the most similar melting curves between replicates most likely have a similar functional state, and thus interactome, between replicates. To generate negative examples, we first manually paired datasets such that each dataset in a pair represented different cell types. We then computed the distance between the melting curve for a given protein between each dataset in a pair for all proteins that were detected in both of the paired datasets. Per dataset, we took proteins whose distance between curves was in the top 80^th^ percentile as true negative examples. This was done with the expectation that proteins with highly different melting curves between different biological contexts most likely have distinct functional states, and thus interactomes. To create training, testing, and validation sets, we took all true positive and true negative proteins and randomly split them first into a train and validation set in an 80:20 ratio. The train set was then further split into a train and test set in an 80:20 ratio. The datasets and dataset pairs were also divided into even sized train and test sets. Final model evaluation was performed using 5-fold cross validation, the results of which are shown in Supplementary Figure S4A.

### Calculating the Intra-Infection Interactome Regulation (IIIR) Score

To calculate intra-infection interactome regulation (IIIR) scores, we first run PROTEA on all possible timepoint combinations (0 and 6, 0 and 12, 12 and 24, etc.). Then, for each timepoint (e.g. 6 HPI) in focus we take the PROTEA score between that timepoint and mock (i.e. 0 HPI), the PROTEA score between the focus timepoint and another infection timepoint (e.g. 12 HPI), and the PROTEA score between this other infection timepoint and mock. Using these three values, we create two points consisting of the infection-infection interactome homology and one of the infection-mock interactome homology (e.g. [6-12 HPI, 6-0 HPI] and [6-12 HPI, 12-0 HPI]). We then compute the distance from this point to the main diagonal using the following formula:

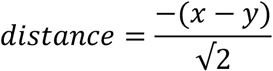

Where *x* = infection-mock interactome homology score and *y* = infection-infection interactome homology score. The sum of these distances is then assigned to the timepoint in focus, and this is repeated for all possible infection-infection pairs that involve the timepoint in focus, generating a list of values. When analyzing temporally resolved interactome regulation, the IIIR score is computed by taking the median value of this list per timepoint. For non-temporally resolved analysis, as was done for generating drug interactome regulation profiles, the max value across all values generated for a given protein within the given infection is used as the infection regulation score. Conceptually, a large positive IIIR score indicates that the interactome of the protein during infection changes a large amount compared to the mock interactome of the protein, and that this change is consistent across infection timepoints, representing a large and sustained interactome change. A large negative IIIR score indicates that there are changes in a proteins interactome between mock and infected states, however these changes are inconsistent across infection timepoints.

### Calculating the Cross-Infection Interactome Regulation (CIIR) Score

To calculate the cross-infection interactome regulation (CIIR) score between for a protein between a pair of infections, we first run PROTEA on paired timepoints between infections. For all comparing amongst measles virus or amongst IAV, HCoV-OC43, DENV, and ZIKV infections all timepoints were paired with themselves (e.g. 0 HPI and 0 HPI, 6 HPI and 6 HPI, etc.). For comparing between measles virus and any other infection the following timepoints were paired: 0 HPI MeV to 0 HPI other virus, 6 HPI MeV to 6 HPI other virus, 12 HPI MeV to 12 HPI other virus, 24 HPI MeV to 18 HPI other virus, 36 HPI MeV to 24 HPI other virus. Using the interactome homology score from each paired timepoint, we calculated the change in interactome homology score between successive timepoints and summed these changes to give the CIIR score. A positive CIIR score indicates that overall, the interactome of a given protein became more similar between experiments as the two infections progressed, while a negative score indicates that the interactome of the protein diverged over the course of the two infections.

### Comparing IIIR and CIIIR scores to Protein Abundance and Stability

#### Protein abundance processing

Protein abundances were calculated using the TPCA reference samples. These samples do not undergo thermal denaturation or separation of soluble and insoluble fractions during the TPCA sample prep. To process these data, the raw protein abundances were first median normalized per sample. Next the resulting values are divided by the median value of the given protein across all timepoints and replicates in which the protein was detected, per protein. These normalization steps are done separately per infection contexts (e.g. independently for MeV infection in MRC5 cells and MeV infection in HFF cells, etc.). Protein abundance fold-changes are then calculated by dividing the mean relative abundance of a protein during an infection timepoint by the mean relative abundance of that same protein of the mock (0 HPI) condition associated with that infection.

#### Protein stability calculation

To calculate the change in the thermal stability of a protein between mock and infected conditions, we utilized the change in protein melting curves between conditions. Specifically, for each protein, we calculate the mean melting curve across replicates per condition. We then calculate the area under the curve for each average curve. The larger the area under the curve is, the more a protein is stabilized against thermal denaturation. To calculate a shift in the relative thermal stability of a protein across biological conditions, we subtract the area under the curve of a protein in the mock condition from its value within an infected condition.

#### Calculating Correlation

To calculate the correlation between the change in abundance of a protein and the interactome of a protein, per infection, we constructed a table of change in protein abundance compared to mock paired with interactome infection regulation scores for all infection timepoints and for all proteins for which both values were available. We then calculated the Pearson’s correlation between resulting list of paired abundance and interactome values. The same methodology same employed to compare protein stability changes to interactome infection regulation, as well as to compare interactome cross-infection regulation to both changes in protein abundance and stability. In the case of comparisons with interactome cross-infection regulation, correlations were calculated per infection pair, instead of per infection.

#### Analysis on the overlap of top regulated proteins and biological processes

To assess the overlap of the topmost regulated proteins as measured by changes in protein abundance, protein stability, and protein interactome (using infection regulation scores), we took filtered proteins to those whose values for these measurements were in the top 95^th^ percentile. Using these sets of proteins, we assessed the overlap in proteins identified by interactome, abundance, and stability per infection. Next, we performed GO term enrichment (using HumanBase^35,138^) on each individual set of proteins and assessed the overlap of biological process GO terms identified by interactome, abundance, and stability.

### Infection UMAPs

In order to construct the infection UMAPs we first constructed a table containing the infection regulation scores at each infected timepoint, the minimum, maximum, mean, and median infection regulation score across all timepoints, the interactome homology scores across all timepoints and possible non-redundant pairs of infections (self-comparisons were assigned NaN values), the cross-infection regulation score across all possible non-redundant pairs of infections (self-comparisons were assigned the value 0), and the minimum, maximum, mean, and median cross-infection regulation score across all infection. All of these values were assembled per protein per infection for the given set of infection being analyzed (i.e. all MeV infections or MeV, IAV, HCoV-OC43, DENV, and ZIKV infections). In the resulting table a row represents a single protein in a single infection, and the columns containing all previously mentioned numeric values, as well as additional columns for tracking protein and infection identity. Next, any row with more than 30 NaN values were removed, followed by imputation of remaining missing values using K-nearest neighbors (KNN) imputation implemented through Scikit-learn python library^139^ with n_neighbors=2000 and weights= ‘distance’. Following imputation, all columns were Z-Score normalized. The resulting Z-Score normalized columns were then used for principal component analysis (PCA) using 14 components. Using these the resulting PCA components, a KNN similarity graph was constructed, using a k of 50 and cosine similarity as the distance metric. The resulting graph was used to perform clustering using the Leiden algorithm. The PCA components were also used to compute a UMAP.

### GO term enrichment

#### Cluster GO term enrichment

To get GO term enrichment of proteins uniquely regulated across a set of infections, the PCA-derived clusters (see Infection UMAPs section of methods) in which the vast majority of proteins within that cluster belonged to the same infection were filtered to in infection regulation score of greater than or equal to 0.08. GO term enrichment was then performed on the resulting set of proteins using HumanBase^138^. To get GO term enrichment of proteins convergently regulated across infections, we combined clusters which showed high average cross-infection regulation scores. We then filtered the resulting set of proteins such that they were present within the combined cluster across a minimum number of infections (5 for MeV infections comparisons, 4 for MeV, IAV, HCoV-OC43, DENV, and ZIKV infections comparison). GO term enrichment was then performed on the resulting set of proteins using HumanBase^138^.

#### Infection pairs GO term enrichment

For a given infection pair, proteins were filtered to the top 90^th^ percentile of cross-infection regulation scores, with the additional requirement of being in the top 50^th^ percentile of infection regulation scores within at least one infection in the pair. GO term enrichment was then performed on the resulting set of proteins using HumanBase^138^.

### SHC1 Interactome PCA

To visualize differences in the temporal regulation of the SHC1 interactome across MeV infection contexts, we constructed a table containing the infection regulation, the interactome homology score within a single infection between a given timepoint and each other timepoint (self-comparisons were assigned a NaN value), and the interactome homology score between each individual infection pair (self-comparisons were assigned a NaN value) at the same timepoint, per timepoint (including 0 HPI and excluding 36 HPI) per infection. For 0 HPI, an infection regulation score of 0 was assigned. KNN imputation was performed to fill in missing values, with n_neighbors=10, and PCA was performed using six components. Per timepoint, the average interactome was computed by calculating the mean point in PCA point using the values of all infections at the given timepoint. The distance between this average interactome and the interactome of each infection was then calculated and Z-score normalized per timepoint.

### Network Representations of Interactomes

In order to aid in the ease of the visualization of protein interactomes in network form, only the top interactions are shown. Specifically, interactions are only shown if they have a Tapioca score greater than 0.5.

### Drug Interactome Regulation Profile Construction

To construct the drug interactome regulation profiles, we first gathered all IIIR and CIIR scores for all protein targets of a given drug. For IIIR scores, per protein target we use the max value across all timepoints of a given infection, we then take the mean of these values across all protein targets of a drug as the IIIR score of the drug. For CIIR scores, we took the mean CIIR scire across all protein targets of the drug. Thus, a drug interactome regulation profile consists of a single IIIR score per infection and a single CIIR per pair of infections.

### Prioritizing of Repurposing Drugs as Antivirals Pipeline

In order to prioritize drugs for potential repurposing as antivirals, we first assembled a set of 7,296 drugs and small molecules with known protein targets from the DrugBank^116^ and ChEMBL^115^ databases. For each of these drugs we constructed drug interactome regulation profiles. Next, we filtered drugs by their infection regulation values, taking only those in the 95^th^ percentile of IIIR scores in at least one infection. In parallel we filtered drugs by their cross-infection regulation values, taking only those in the 98^th^ percentile of CIIR scores in at least one infection pair. We then took forward only the drugs that passed both of these filters. Next, we used top GO terms identified from our previous GO term analysis of convergent biological processes to filter to drugs whose protein targets are known to be involved in at least one of these biological processes as annotated by UniProt^140^. In the next step of filter, we calculated the max IIIR and CIIR scores across all infections and infection pairs per drug. Using these values, we ranked the remaining drugs separately by IIIR and CIIR scores, taking forward the drugs that were either in the top 60 or 90 of each ranked listed respectively. Next, we filtered the remaining drugs to those with a maximum of 10 known protein targets, and for whom at least one third of their known protein targets were detected across all infections, giving a final set of 92 prioritized drugs and small molecules.

### Comparing Protein Regulation Between All MeV and Non-MeV Infections

Each non-MeV infection (i.e. HCoV-OC43, IAV, DENV, ZIKV) was individually compared to each MeV infection across all cell types. First, for each non-MeV infection, proteins within the top 90^th^ percentile of IIIR scores, protein abundance log fold change, and protein thermal stability shift (all absolute value), were selected for each infection, forming sets of top regulated proteins by interactome, abundance, and stability, respectively. For each non-MeV and MeV infection pair, the CIIR score and protein abundance and protein thermal stability correlation was calculated for all proteins identified as amongst the most regulated within the non-MeV infection. The distribution of these scores was then compared between non-MeV infections, both resolved by MeV infection and grouped, to identify which non-MeV infections converged most with MeV infections by interactome, abundance, or stability changes.

### TPCA MS Sample Preparation

#### Collection and denaturation

Cells were collected via trypsinization and then resuspended in 1x PBS, after which 50 μl of cells were aliquoted into PCR strips, eight tubes for denaturation and one additional tube for a non-denatured reference sample (i.e. another sample of cells from the same culture). Denaturation was then immediately performed on the relevant samples. Thermal denaturation for TPCA samples was performed using 8 temperature points between 37 °C and 55 °C (37 °C, 38.2 °C, 40.3 °C, 43.9 °C, 48.1 °C, 51.2 °C, 53.5 °C, 55 °C). After denaturation, 1.5x kinase buffer (75 mM HEPES [Sigma Aldrich, H4034-500G] pH 7.5, 15 mM MgCl_2_ [ThermoFisher Scientific, 447155000], 1.5× Halt protease and phosphatase inhibitor [ThermoFisher Scientific, 78438], 3 mM tris(2-carboxyethylphosphone) [TCEP, ThermoFisher Scientific, 77720]) were added to all samples, including reference samples. Samples were then snap-frozen and stored at -80 °C until preparation for MS analysis.

#### MS sample preparation

Denatured samples were thawed on ice and then lysed in 1% Triton X-100 (ThermoFisher Scientific, A16046.AE), 0.1% Tween-20 (Sigma Aldrich, P1379-100ML), and 0.5% sodium deoxycholate (Neta Scientific, SIAL-D6750-25G) in 20 mM HEPES, 110 mM KOAc (ThermoFisher Scientific, AC418175000), 2 mM MgCl_2_, 1 μM ZnCl_2_ (ThermoFisher Scientific, A16281.36), and 1 μM CaCl_2_ (ThermoFisher Scientific, 423525000) at pH 7.4 for 1 h. The samples were then further lysed via three freeze-thaw cycles. Reference samples were lysed in 5% SDS (Sigma-Aldrich, L4509-10G) for 1 h. Denatured samples were then centrifuged at 20,000g for 20 min at 4 °C, pelleting the insoluble protein. The soluble proteins in the supernatant were then transferred to a new 1.5 ml low-bind tube. The remaining insoluble protein pellet was resuspended in 10% SDS. All samples were then reduced and alkylated with 25 mM TCEP and 25 mM chloroacetamide (Neta Scientific, SIAL-22790-250G-F) at 55 °C for 30 min. Reduced and alkylated samples were then purified via methanol-chloroform precipitation. The resulting protein disks were then resuspended in 100 mM HEPES (pH 8.3). Following determination of sample protein concentration by BCA assay, samples were aliquoted into new low-bind tubes at a volume giving a concentration of 0.5 mg/ml for the 37 °C. The same volume used at 37 °C was used for all other temperatures within a given set. The aliquoted samples were then digested at 37 °C with 1 μg of sequencing grade trypsin (ThermoFisher Scientific, PI90059) for 16 h. Trifluoroacetic acid (TFA, ThermoFisher Scientific, PI28904) was then added to digested samples to a final concentration of 1% TFA. Stage tips were prepared containing disks of SDB-RPS (CDS Analytical, 98-0604-0226-4EA). The stage tips were first washed with methanol, followed by a wash with buffer B (0.1% formic acid [FA, ThermoFisher Scientific, 28905], 80% acetonitrile [ACN, ThermoFisher Scientific, A9554]), then two washes with buffer A (0.1% FA). Samples were then spun at 20,000g for 5 min to pellet any insoluble containments. Sample supernatant was then passed through stage tips, one sample per stage tip. Stage tips were then washed once with buffer A and twice with buffer B. Peptides were eluted using elution buffer (5% ammonium hydroxide, 80% ACN). Peptides were then dried down by speed vacuum centrifugation and then resuspended in 0.1% FA and 4% ACN, prior to analysis on a timsTOF Ultra (Bruker). Samples were resuspended such that all samples were injected at equal protein concentration.

#### Peptide liquid chromatography-tandem MS

LC-MS/MS analysis was performed with a nanoElute2 (Bruker) coupled to a timsTOF Ultra (Bruker) with 1 μl injections (150 ng of peptide on column). The mobile phases were 0.1% FA in 99.9% UHPLC-MS water (ThermoFisher Scientific, W81CS, buffer A) and 0.1% FA in 99.9% UHPLC-MS acetonitrile (ThermoFisher Scientific, A9561CS, buffer B). A one-column separation method was used with a PepSep ULTRA (250 mm × 75 μm x 1.5 μm) C18 HPLC column (Bruker ,1893484) as the analytical column. A 10 μm emitter (Bruker, 1811112) attached to a CaptiveSpray Ultra source with a column toaster set to 50 °C was used. A linear 30-min gradient of 3% to 34% buffer B at a flow rate of 200 nl/min was used for peptide separation.

#### LC-MS/MS analysis

For data-independent acquisition parallel accumulation serial fragmentation (dia-PASEF) analysis, the MS1 settings were set to start at 100 m/z and end at 1700 m/z in positive ion mode. For the TIMS settings, the mode was set to custom, with a starting 1/K0 of 0.65 V s/cm2 and an ending 1/K0 of 1.46 V s/cm2. Collision energy was linearly scaled to the ion mobility, ranging from 20 eV to 59 eV between 0.65 V s/cm2 and 1.46 V s/cm2. The ramp time was set to 50 ms with a 100% duty cycle and a ramp rate of 17.80 Hz. A 16 x 3 method was used for DIA windows, three groups of 16, spanning the mobility and m/z ranges. The estimated cycle time was 0.95 s.

#### Peptide identification and quantification

DIA data were analyzed with DIA-NN^141^ (version 1.9), where a search was performed using a merged database of Homo sapiens appended with common contaminates (downloaded 02/05/2024 from Bruker) and viral proteins. A separate merged database was made per virus using the following proteomes: UP000008699 (MeV), downloaded from UniProt on 07/31/2024 UP000009255 (IAV) downloaded from UniProt on 02/01/2024, UP000007552 (HCoV-OC43) downloaded from UniProt on 07/31/2024, UP000007196 (DENV) downloaded from UniProt on 08/16/2024, and UP000054557 (ZIKV) downloaded from UniProt on 08/16/2024. Both the DENV and ZIKV proteomes on UniProt are as a single polypeptide. Therefore, both UP000007196 and UP000054557 were further processed into separate viral proteins based on the annotations within the polypeptide UniProt entries (UniProt entries: P14340, DENV; Q32ZE1, ZIKV). These cleaved proteomes were then used for peptide identification. A spectral library was produced with this database via DIA-NN, where the fragment ion m/z range was set to 300-1300 m/z, N-terminal methionine excision was enable, in silico digestion with trypsin with cleavage at lysine and arginine, a maximum of one missed cleavage, a peptide length between 7 and 30 amino acids, a precursor m/z range of 100 to 1,700, precursor charge range of 1 to 4, a fixed cysteine carbamidomethylation, with no dynamic modifications. Precursor ion and fragment ion mass tolerance were both set to 0.0 to allow DIA-NN to determine the appropriate cutoffs. For DIA-NN search settings, unrelated runs, MBR, and no shared spectra were enabled. The neural network classifier was set to single-pass mode, the quantification strategy was set to QuantUMS (high precision), and cross-run normalization was set to off. Peptides and proteins filtered at a 1% FDR were used for downstream analysis.

#### Processing of raw TPCA data

First, using the pr_matrix and pg_matrix files outputted by DIA-NN, proteins with less than two identified proteins were discarded. Next, a per sample (one temperature of one condition of one replicate) a scaling factor was computed by dividing the MS2.Signal value (found in the .stats file outputted by DIA-NN) associated with that sample by the mean MS2.Signal value across all analyzed samples. All abundance values with a sample were then multiplied by their respective sample-specific scaling factor. Next, for each protein, its reference abundance, specific to both condition and replicate, but not temperature, was divided by the median reference abundance for the given protein across all conditions and replicates. All abundance values for a given protein within a given condition and replicate were then multiplied by their respective condition and replicate specific reference scaling factor. For samples in which the protein was detected in the thermally denatured samples, but not the reference sample, all abundance values for a given protein within the given condition and replicate were multiplied by the median reference scaling factor respective to the given condition and replicate. Next, all abundances for a given temperature were multiplied by that temperatures associated volume scaling factor to account for equalization of protein concentration across samples prior to MS analysis (37 °C: 1.0, 38.2 °C: 1.0, 40.3 °C: 1.0, 43.9 °C: 1.0, 48.1 °C: 0.94, 51.2 °C: 0.73, 53.5 °C: 0.6, 55 °C: 0.5). Median of median normalization was performed next. First, the median abundance value across all proteins within each individual sample (one temperature of one condition of one replicate) was calculated. The median value of all of these median values was then calculated, and then all abundances were divided by this median value. Following this, for all proteins, per condition, the median abundance of a given protein across all temperatures and replicates was calculated. All abundance values for that protein within the specific condition were then divided by this median value. Next, proteins whose curves within a given condition and replicate contained any missing values were discarded. Finally, the data were fit to a three-parameter log-logistic equation:

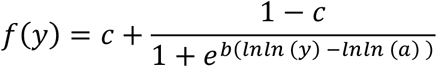

### PPI Prediction from TPCA Data

PPIs were predicted from TPCA data using Tapioca^23^, a machine learning-based framework previously developed by our group that integrates TPCA data with protein physical properties, protein domains, and tissue-specific functional networks. Here, a skin-specific functional network was used for data collected in HFF and A375 cells, a lung-specific functional network was used for data collected in MRC5 and A549 cells, an eye-specific functional network was used for data collected in ARPE19 cells, a neuron-specific functional network was used for data collected in BE(2)-C cells, a monocyte-specific functional network was used for data collected in THP-1 cells, and a hepatocyte-specific functional network was used for data collected in HuH7 cells.

### Cell Lines and Primary Cell Culture

A375 (ATCC, CRL-1619), A549 cells (ATCC, CCL-185), HFF cells (ATCC, SCRC-1041), HuH-7 cells (a gift from A. Ploss, Princeton University), MRC5 cells (ATCC, CCL-171), Vero (a gift from Neal DeLuca, University of Pittsburgh), were cultured in DMEM media (ThermoFisher Scientific, 21013024). ARPE-19 cells (ATCC, CRL-2302) were grown in DMEM:F12 media (ThermoFisher Scientific, 12634010), BE(2)-C cells (ATCC, CRL-2268) were grown in EMEM:F12 media (Millipore Sigma, SLM-246), and THP-1 cells (ATCC, TIB-202) were grown in RPMI-1640 media (Millipore Sigma, R8758). All media was supplemented with 10% fetal bovine serum (FBS, Gemini Bio, 100-106-500) and 1% penicillin-streptomycin (Millipore Sigma, P4458) unless otherwise stated. All cells were grown at 37°C and 5% CO2 unless otherwise stated.

### Virus Strains and Propagation

Measles virus Edmonston strain (ATCC, VR-24) was propagated in Vero cells. After acquisition of the virus, it was propagated once to produce a P0 stock, which was then used to produce P1 stocks which were used for all experimentation. For both P0 and P1 stocks, Vero cells were infected at a low multiplicity of infection (MOI) and virus was collected once greater than 90% of cells showed cytopathic effect. Virus was collected through scrapping cells and collecting both the cells and infected media. These samples were then gently sonicated to release virus from cells before being centrifuged at 1,000g for 10 minutes to pelleted cell debris. The supernatant was then aliquoted and stored at -80 °C until use. The virus was titered in using immunofluorescent imaging using a Measles virus monoclonal antibody (ThermoFisher Scientific, MA5-47564) to detect infected cells. The virus was titered separately for each cell line in which an infection was performed. Human coronavirus OC43 (gift from Dr. Kevin Harrod, University of Alabama at Birmingham; originally purchased from ATCC, VR-1558) was propagated in MRC5 cells at a low MOI, collecting virus once greater than 90% of cells showed cytopathic effect, to produce a P1 stock. Virus was collected in the same manner as the measles virus stock. The virus was titered in MRC5 cells using immunofluorescent imaging using Anti-Coronavirus Group Antigen Antibody, nucleoprotein of OC-43, clone 542-7D (Millipore Sigma, MAB9013). Influenza A virus H1N1 A/Puerto Rico/8/1934 strain (a gift from Dr Thomas Shenk, Princeton University) was propagated in A549 cells. As part of a prior study by our group ^50^, this virus had been serially propagated for 10 times in A549 cells to improve virus yield for TPCA experiments and virus stocks were stored at -80 °C until use. Virus was collected in the same manner as previously described above. In this study, the virus was titered in MRC5 cells using immunofluorescent imaging using an Anti-Influenza A Virus Nucleoprotein antibody (abcam, ab128193) to detect infected cells. Dengue virus type 2 (ATCC, VR-1584) and Zika virus PRVABC59 strain (ATCC, VR-1843) were propagated at a low MOI in Vero cells, collecting virus once greater than 90% of cells showed cytopathic effect, to produce a P1 stock. Virus was collected in the same manner as previously described above. These viruses were titered via TCID50 in HuH-7 cells.

### Viral Infections for TPCA Experiments

In TPCA experiments viral infections were performed at a MOI of 3 for measles virus, a MOI of 3 for human coronavirus OC43, a MOI of 5 for influenza A virus, a MOI of 10 for dengue virus, and a MOI of 10 for Zika virus. For a one-hour incubation period cells were infected in half-volume of their respective growth media, with 0% FBS. In the case of the influenza A infection, this media was supplemented with 0.25 µg/ml of TPCK trypsin (ThermoFisher Scientific, 20233). Following incubation, the infected media was removed, cells were washed with warm PBS, and then the full volume of their respective growth medias, complete with 10% FBS, were added back to the cells. When working with THP-1 cells, at any step where media was removed the cells were first centrifuged at 500g for 10 minutes to pellet cells. Infected cells were harvested at the following hours post infection (HPI) for each virus: 0, 6, 12, 24, and 35 HPI for all measles virus infections, and 0, 6, 12, 18, and 24 HPI for all human coronavirus OC43, influenza A virus, dengue virus, and Zika virus infections. In the case of 0 HPI timepoints, no virus was added to the cells, however they were otherwise handled in the same manner as all infected timepoints, in order to serve as appropriate controls. These experiments were conducted in biological replicates of three.

### Drug Treatment Infections & Titering

For drug repurposing experiments, cells were infected in the same manner was done for TPCA up to the end of the 1-hour infection incubation period, with the exception that all infections were done at an MOI of 0.01. Following the incubation period, the infected media was removed, cells were washed with warm PBS, and then growth media containing the experimental drugs and 0% FBS were added to the cells. Drugs were used at the following final concentrations: 49.40 nM voclosporin (Neta Scientific, CAYM-39719-1), 133.00 µM etidronic acid (ThermoFisher Scientific, 501970105), 1.97 µM clodronic acid (used disodium clodronate tetrahydrate, ThermoFisher Scientific, 502192270), 200.00 nM auranofin (Neta Scientific, CAYM-15316-25), 3.00 nM ixabepilone (Neta Scientific, CAYM-23732-1), 5.23 nM sacubitril (used sacubitril calcium salt, Sigma-Aldrich, SML1380-5MG), 6.20 nM nirogacestat (ThermoFisher Scientific, 501873146), 30.00 µM pyrimethamine (Neta Scientific, CAYM-16472-10), 30.00 µM emapunil (ThermoFisher Scientific, 501873270), 350.00 nM BMS-309403 (ThermoFisher Scientific, 502054013). Etidronic acid and clodronic acid were dissolved in autoclaved water before being added to growth media. The remaining drugs were dissolved in DMSO (Neta Scientific, SIAL-D4540-100ML) before being added to growth media, and the final concentration of DMSO in the media was 0.1% DMSO. All drugs were sterile filtered after being dissolved and before being added to the media. Sterile filtered water and 0.1% DMSO were used as controls. After drugged media was added to cells, the infection was allowed to progress. To maintain drug concentration over the time course of the experiment, drugs were spiked in at half final concentration at 24 HPI, and at full final concentration at 48 HPI. Once the control samples showed greater than 90% cytopathic effect, determined individually per infection, virus from all control and drug samples related to that infection were collected following the same methodology as described for viral propagation. The maximum time to 90% cytopathic effect observed across all experiments was 72 hours. All virus was titered using TCID50 in Vero cells for measles, dengue, and Zika infections, and in MRC5 cells for influenza A and human coronavirus OC43 infections. These experiments were conducted in biological replicates of three.

### Drug Treatment Control Experiments

For drug treatment control experiments, cells were treated in the exact same manner as in drug treatment infection experiments, except for no viral infection occurred and the all drug treatment went to the full 72 hours, which was the maximum time to 90% cytopathic effect observed for drug treatment infection experiments. At 72 hours, media was removed and cells were washed with warm PBS before being trypsinized. After cells had de-adhered, trypan blue solution (ThermoFisher Scientific, 15250061) was added to the cell suspension at a to 1:1 ratio. The percentage of alive cells were then recoded using a Countess II automated cell counter (ThermoFisher Scientific). These experiments were conducted in biological replicates of three.

### Quantification and Statistical Analysis

Data processing and large-scale analyses were performed using Python^135^ utilizing the Python libraries Numpy^142^, Scipy^143^, Pandas^144^, Networkx^145^ , igraph^146^, Scikit-learn^139^, Statsmodels^147^, UMAP-learn^148^, Seaborn^149^, and Matplotlib^150^. For calculating statistics, the method to calculate p-values and for FDR correction of p-values are mentioned in the relevant figure caption. For all experiments three biological replicates were performed. Some network visualizations were generated using CytoScape^151^ . Figures were created using Python and Microsoft Powerpoint.

