## Supplemental Figures for "A Temporal Interactome Atlas across RNA Viruses reveals Convergent Vulnerabilities for Antiviral Repurposing"

###### **This Supplementary Information Contains:**

Supplementary Figures S1 to S12

Supplemental References

##### Number of Proteins With Abundance Measurements Across Infections

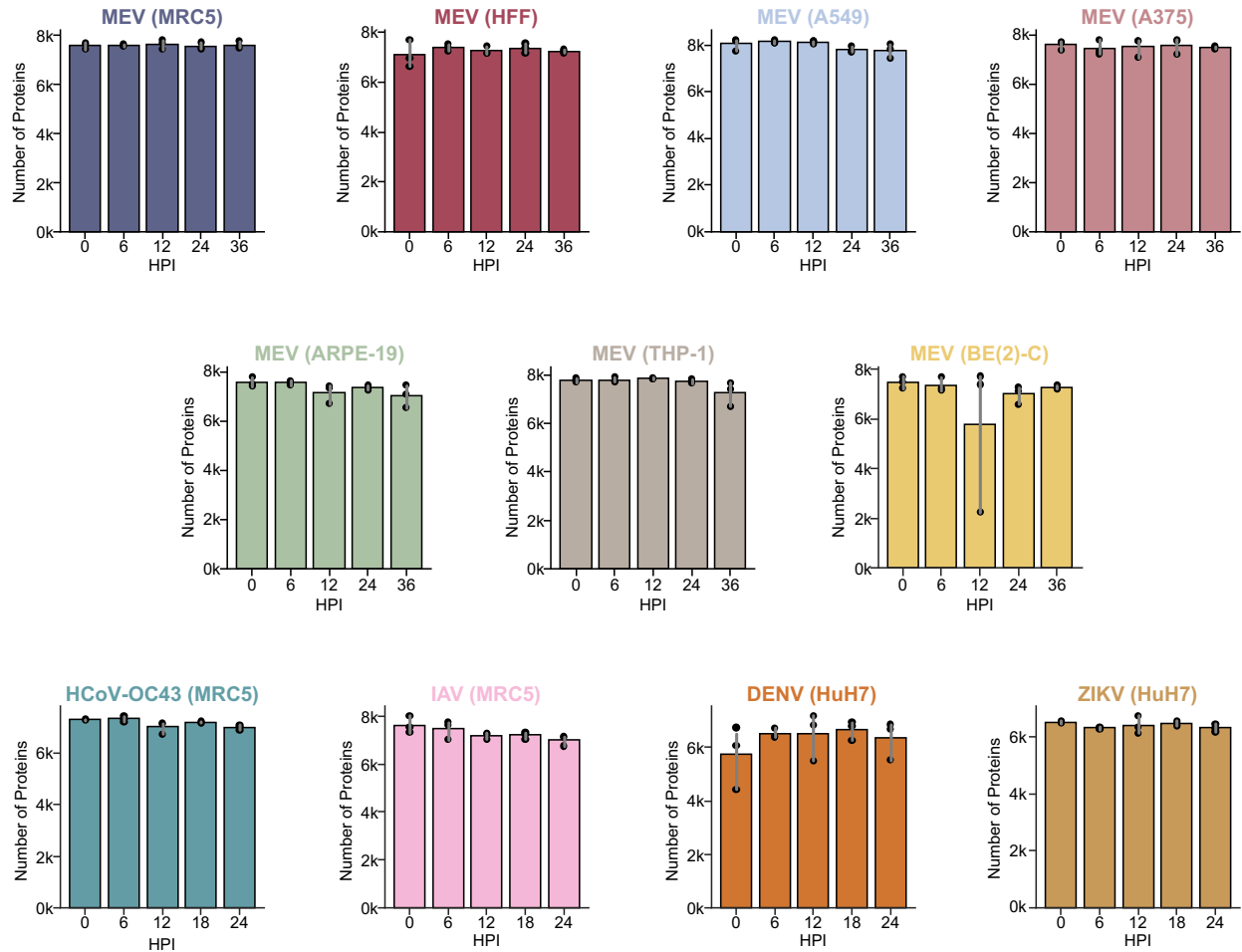

**Figure S1. Overview of protein abundance data in the Interactome & Viral Infection Dynamics (InterVir) Atlas.** Bar charts showing the average number of protein abundance data points per timepoint and infection. In the plot, the grey line represents the 95% confidence interval, and the black dots show the exact value per each of the three biological replicates.

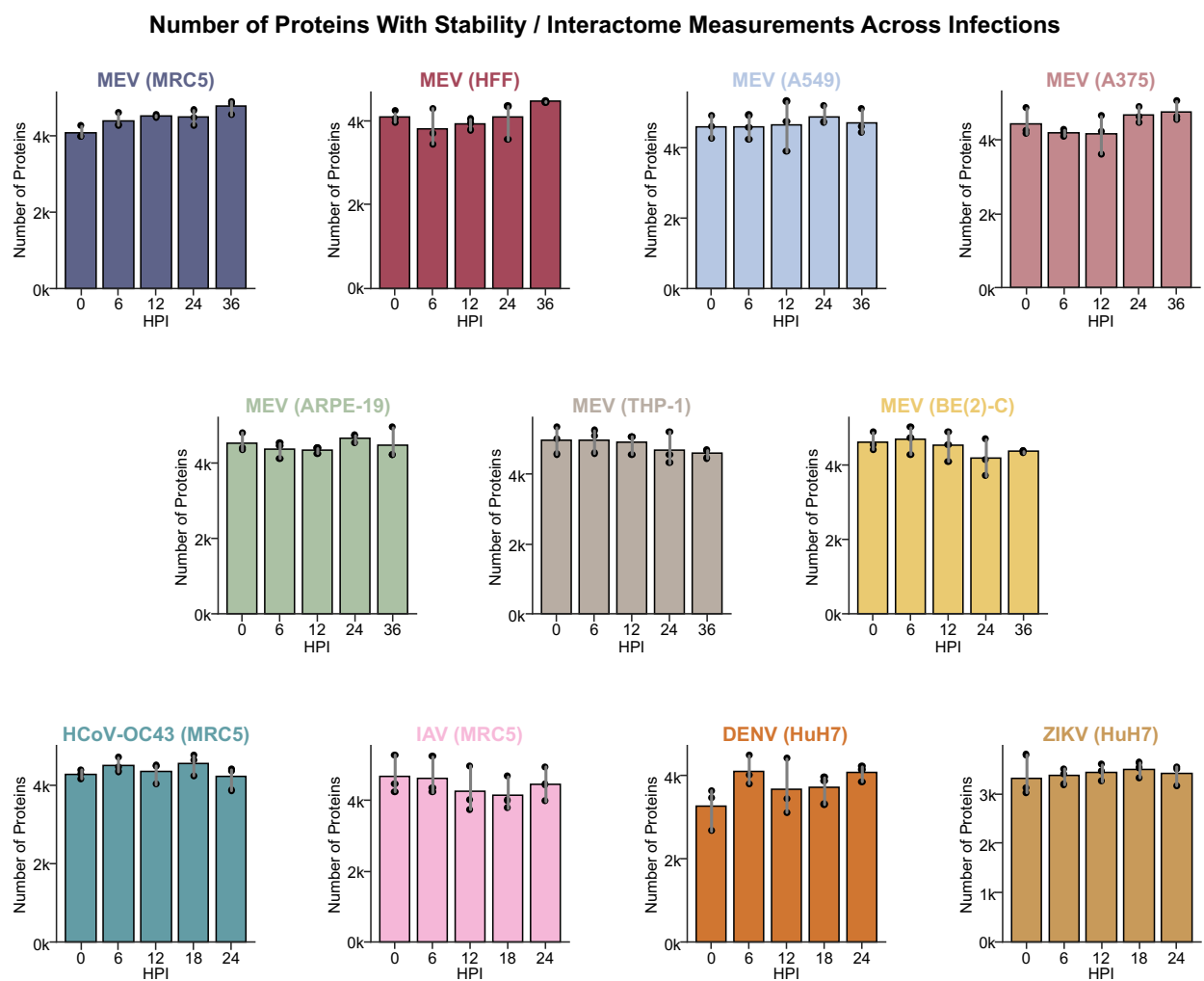

**Figure S2. Overview of protein stability and interactome data in the InterVir Atlas.** Bar charts showing the average number of protein stability and data points per timepoint and infection (one protein stability point equals one complete protein melting curve equals one protein interactome). In the plot, the grey line represents the 95% confidence interval, and the black dots show the exact value per each of the three biological replicates.

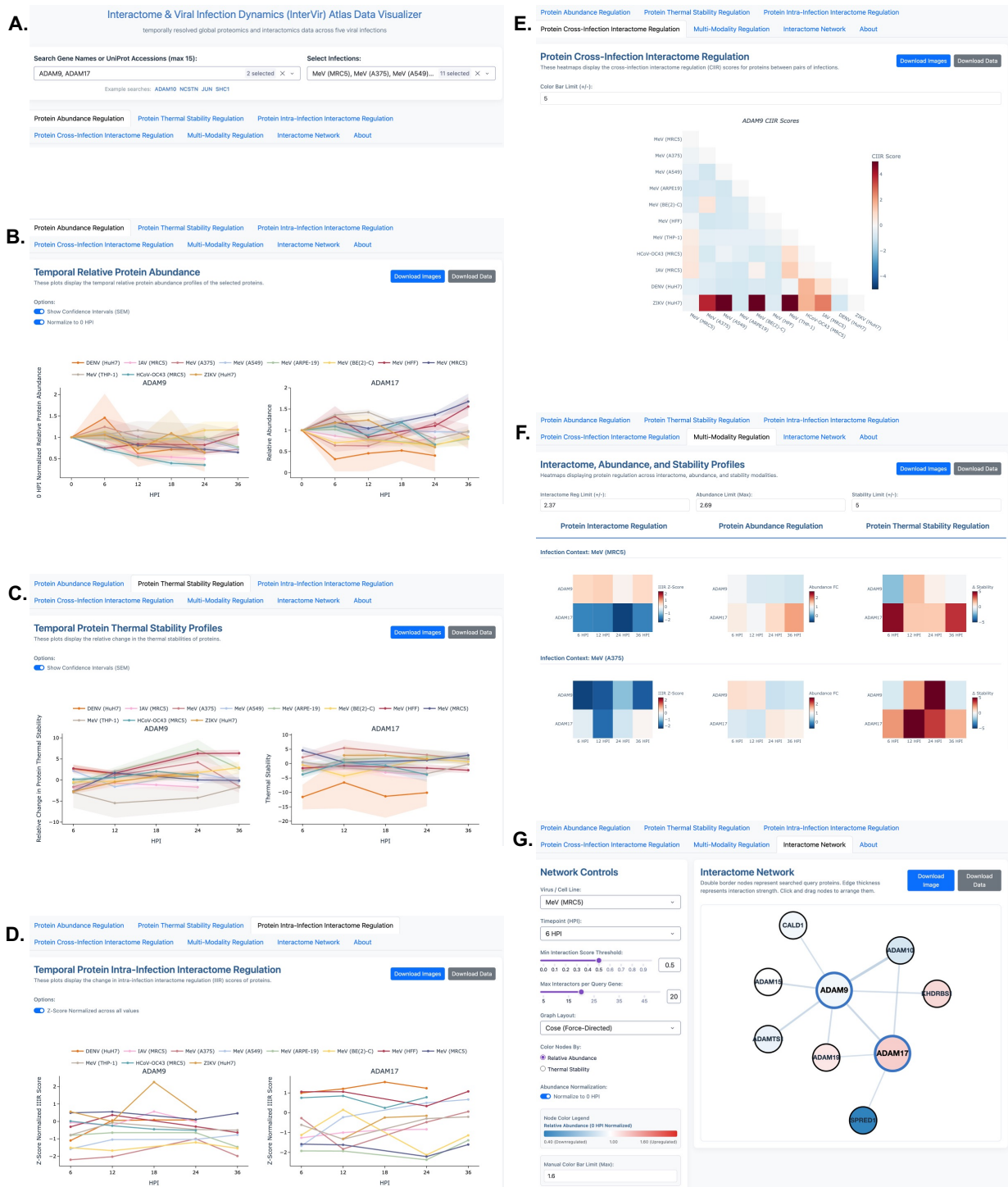

**Figure S3. Overview of the InterVir Atlas data visualization website.** We developed a data visualization website (https://interviratlas.princeton.edu/) to enable accessible exploration of our InterVir Atlas. For all plots and networks, the image and underlying data can be downloaded for further customized analysis. **A**, Users can search for proteins by gene name or uniprot accession and select multiple proteins to visualize (up to 15 at once). Users can also search and select to visualize up to all of the infection contexts in the InterVir Atlas. **B**, Line plots of temporal protein

abundance can be generated, showing the abundance across all select infections and generating separate plots per selected proteins. **C**, Line plots of temporal protein thermal stability can be generated, showing the stability across all select infections and generating separate plots per selected proteins. **D**, Line plots of temporal protein intra-interaction interactome regulation (IIIR) scores can be generated, showing the IIIR scores across all select infections and generating separate plots per selected proteins. **E**, Heatmaps can be generated visualizing the cross-infection interactome regulation (CIIR) scores across all pairs of selected infections. A separate heatmap is generated per each selected protein. **F**, To compare regulation of across protein abundance, thermal stability, and IIIR scores across multiple proteins within an infection, users can generate multiple heatmaps showing temporal values for all selected proteins. A separate heatmap is generated per data modality and infection. **G**, A network visualization of the interactome(s) of the selected proteins can be generated for a given infection and hour post infection (HPI). The score threshold and maximum number of interactors per protein can be adjusted. Nodes can be colored by relative change in protein abundance or thermal stability compared to 0 HPI.

##### A. Protea Performance

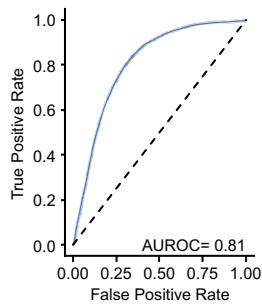

### B.

##### Interactome Regulation Does Not Correlate with Protein Abundance or Stability Regulation

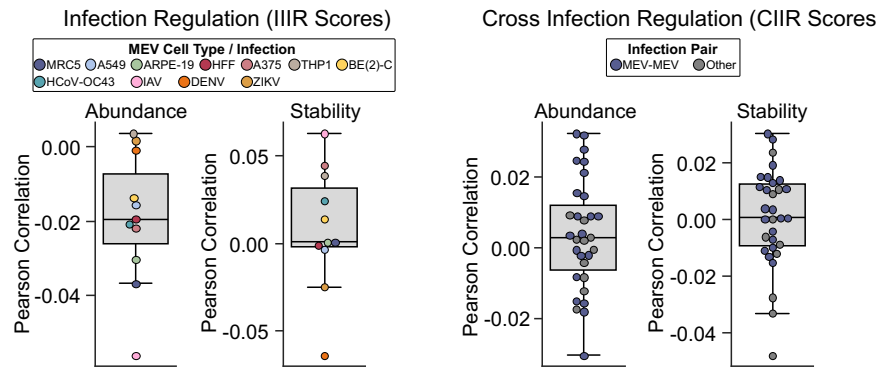

**Figure S4. Protea performance and the correlation between CIIR scores and protein abundance and stability.** **A**, An area under the receiver operating characteristic curve (AUROC) showing Protea performance in 5-fold cross validation (see methods for details). **B**, Box plots showing the correlation between IIIR or CIIR scores and changes in protein abundance or stability across viral infections. The line within the box represents the median value and the whiskers represent the $\pm 1.5$ interquartile range.

#### GO Term Enrichment of MEV Infection Interactome Regulation Biological Processes

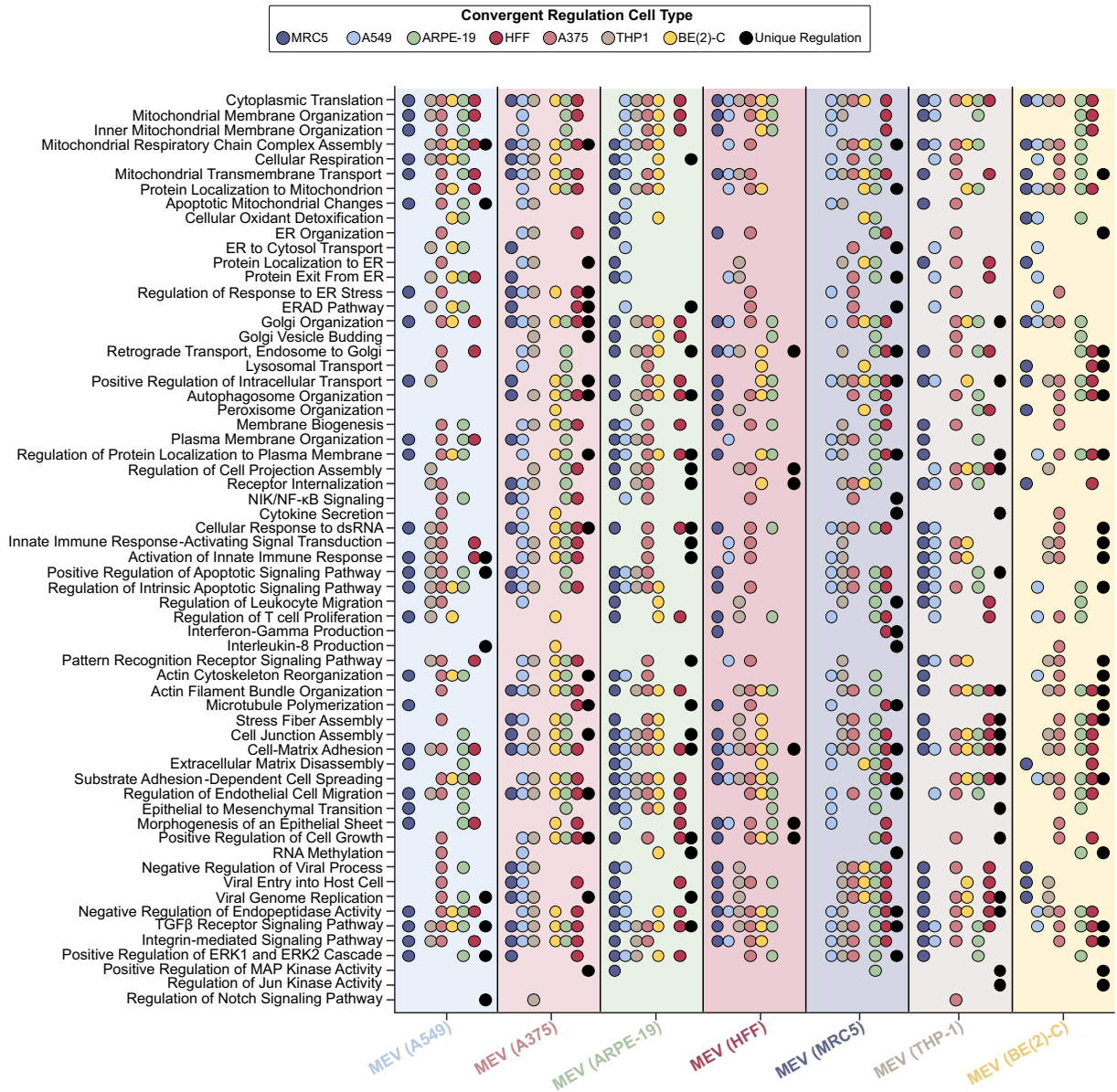

**Figure S5. GO term enrichment of convergently and divergently regulated protein interactomes across MeV infections in different cell types.** GO term enrichment of proteins whose interactomes were convergently regulated during MeV infection between a pair of cell lines (colored dots) or were uniquely regulated within a given cell line (black dots). GO term enrichment was performed using HumanBase<sup>1</sup>.

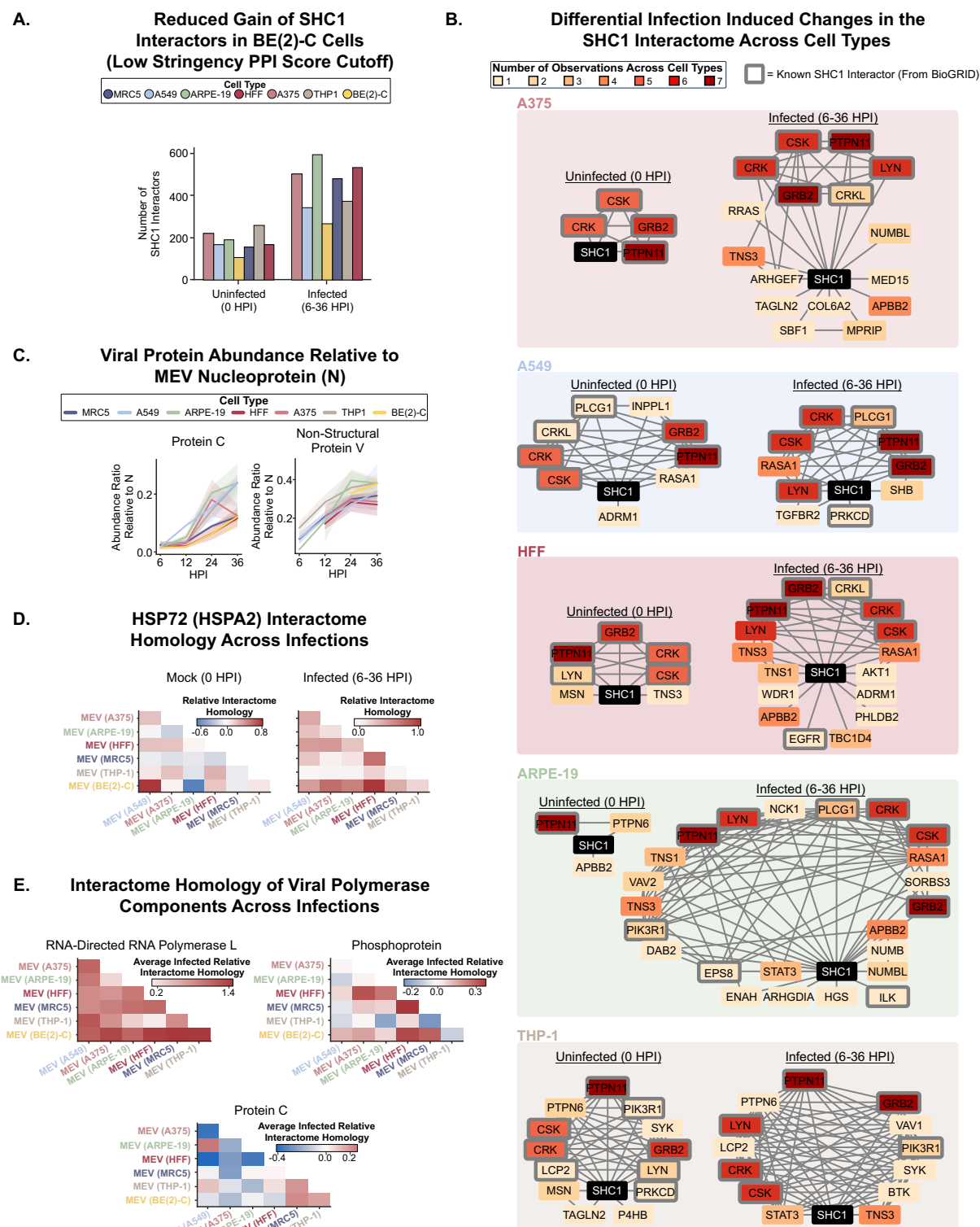

**Figure S6. Regulation of the interactomes and abundances of host and viral proteins across MeV infections in different cell types. A,** Bar plot depicting the number of SHC1 interactors (including low-confidence predictions) at uninfected and infected states per cell line. **B,** SHC1

interactomes during uninfected and infected states. **C**, Temporal abundances of select viral proteins per cell line normalized to respective viral nucleoprotein abundances. The solid line represents the median value, and the shaded region represents the 95% confidence interval. **D**, Interactome homology of HSP72 (HSPA2) between all cell line pairs used for MeV experiments at both mock (0 HPI) and averaged across all infected timepoints (6-36 HPI). **E**, Average interactome homology values of viral protein members of the viral polymerase complex compared between all measured MeV infections.

### GO Term Enrichment Biological Processes With Interactome Regulation Across MEV, HCoV-OC43, IAV, DENV, and ZIKV Infections

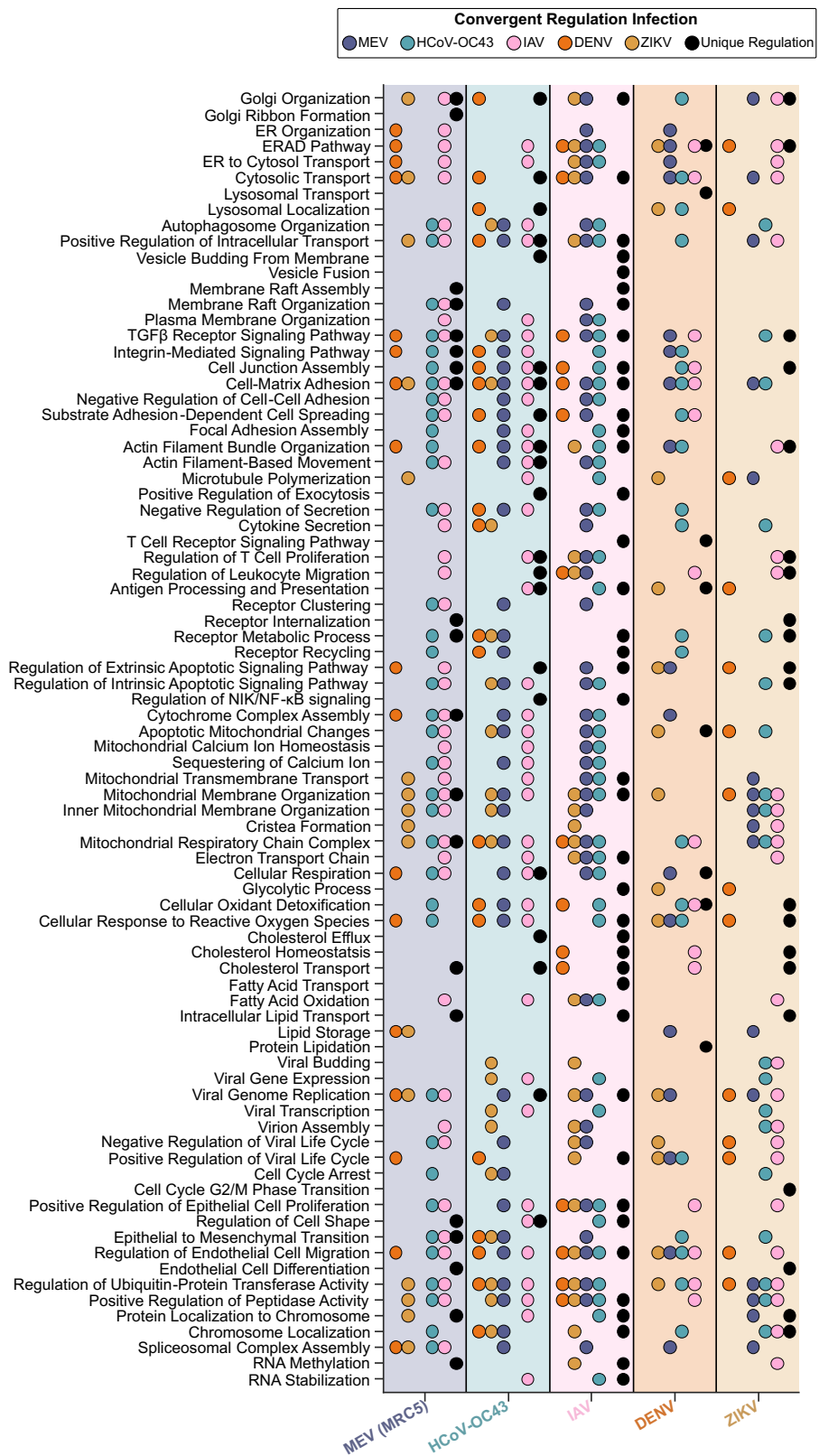

**Figure S7. GO term enrichment of convergently and divergently regulated protein interactomes measles virus (MeV), human coronavirus OC43 (HCoV-OC43), influenza A (IAV), dengue (DENV) and Zika (ZIKV) infections. A, GO term enrichment of proteins whose interactomes were convergently regulated during infection between a pair of viruses (colored dots) or were uniquely regulated within a virus (black dots). GO term enrichment was performed using HumanBase<sup>1</sup>.**

### A. Cross Infection Regulation of Immune Related Protein Interactomes

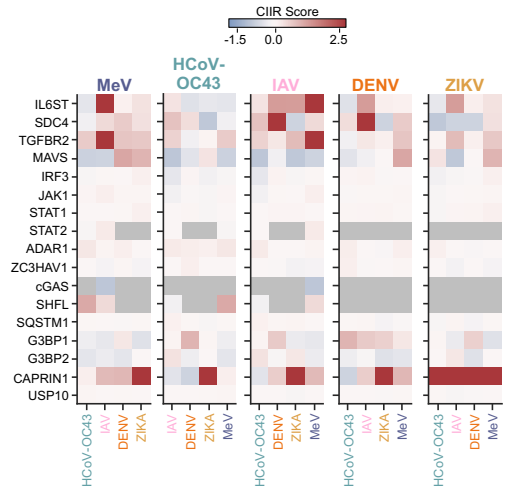

### B. Changes in Immune Related Protein Relative Abundance

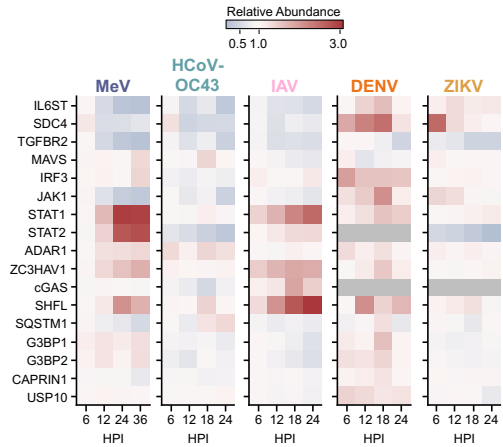

### C. Regulation of Immune Related Protein Thermal Stability

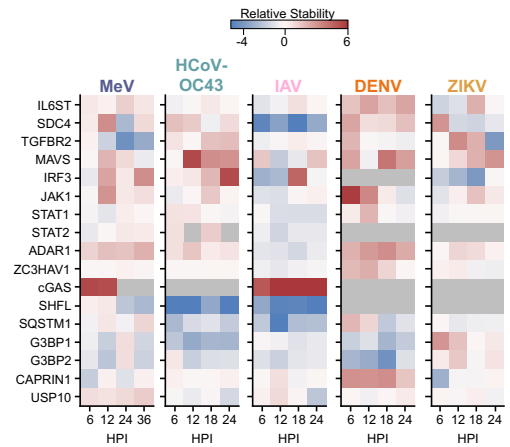

**Figure S8. Interactome, abundance, and stability regulation of immune proteins across different viral infections.** **A**, Heatmap showing the CIIR scores of immune proteins across different sets of infection comparisons. The infection at the top each individual heatmap is the infection in focus, and the infection at below column represents the other infection being compared to the infection in focus. **B**, Heatmap showing the relative protein abundances of immune proteins across each timepoint of infection for each infection. Protein abundances are normalized to the 0 HPI abundance. **C**, Heatmap showing the relative protein thermal stability of immune proteins across each timepoint of infection for each infection. Protein thermal stabilities have been Z-score normalized.

**A. Cross Infection Regulation of Membrane Contact Site Protein Interactomes**

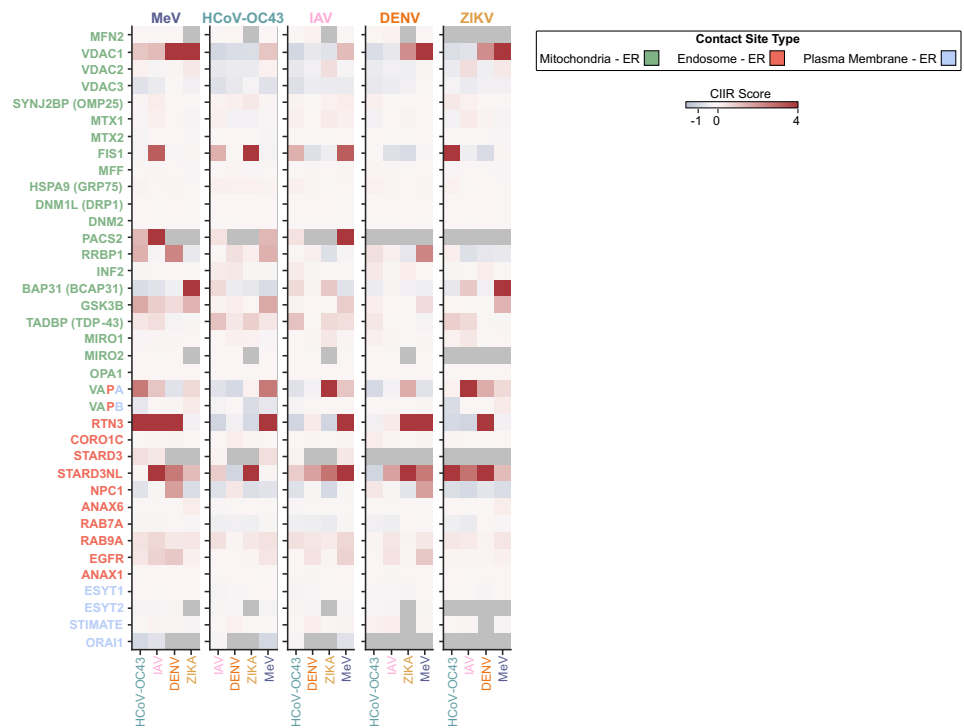

**B. Changes in Membrane Contact Site Protein Relative Abundance**

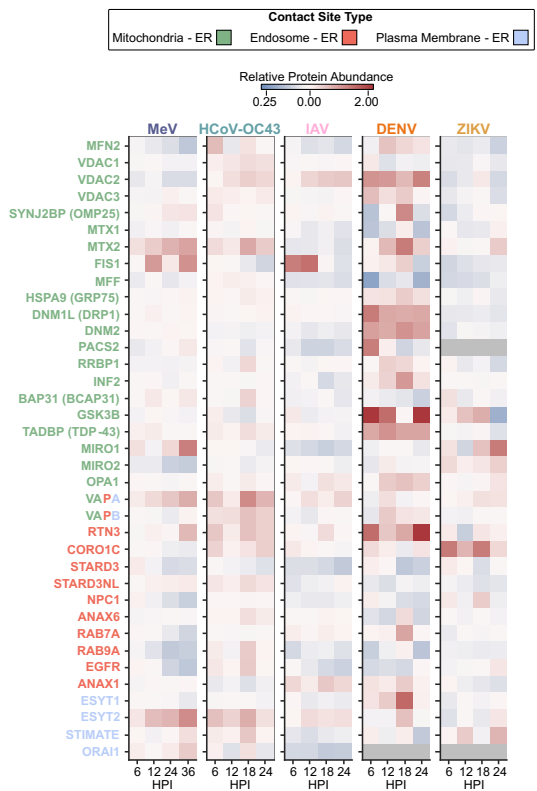

**C. Regulation of Membrane Contact Site Protein Thermal Stability**

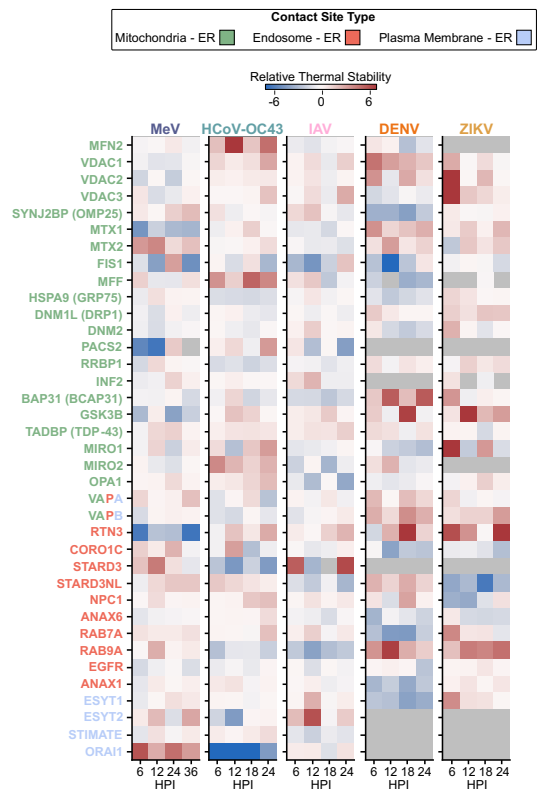

**Figure S9. Interactome, abundance, and stability regulation of membrane contact site (MCS) proteins across different viral infections.** **A**, Heatmap showing the CIIR scores of MCS proteins across different sets of infection comparisons. The infection at the top each individual heatmap is the infection in focus, and the infection at below column represents the other infection being compared to the infection in focus. **B**, Heatmap showing the relative protein abundances of MCS proteins across each timepoint of infection for each infection. Protein abundances are normalized to the 0 HPI abundance. **C**, Heatmap showing the relative protein thermal stability of MCS proteins across each timepoint of infection for each infection. Protein thermal stabilities have been Z-score normalized.

#### Alpha Secretases Display a Unique Interactome in DENV Infection

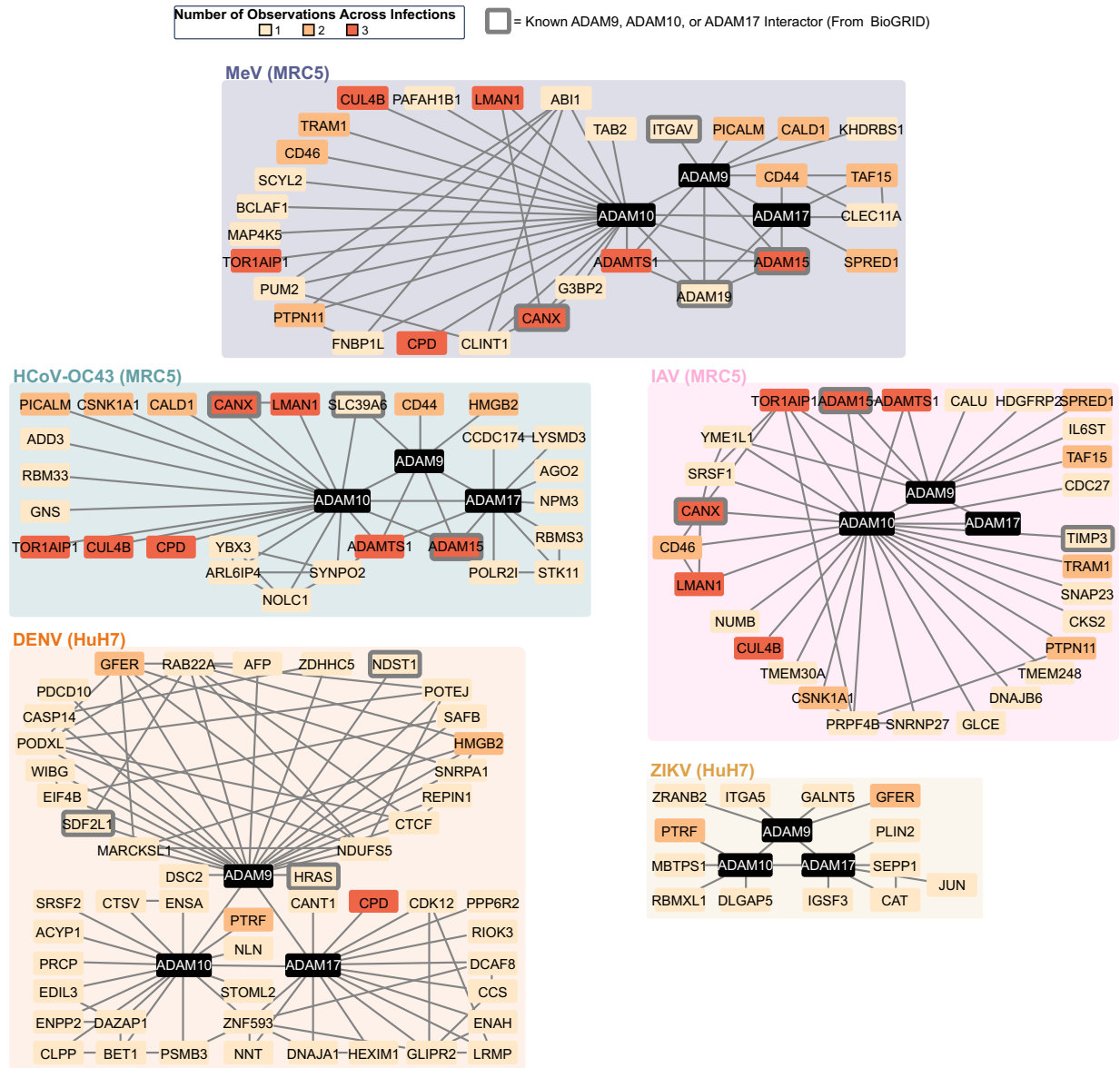

**Figure S10. Interactomes of alpha secretases during infection with different viruses.** Alpha secretase interactomes during infection with MeV, HCoV-OC43, IAV, DENV, or ZIKV.

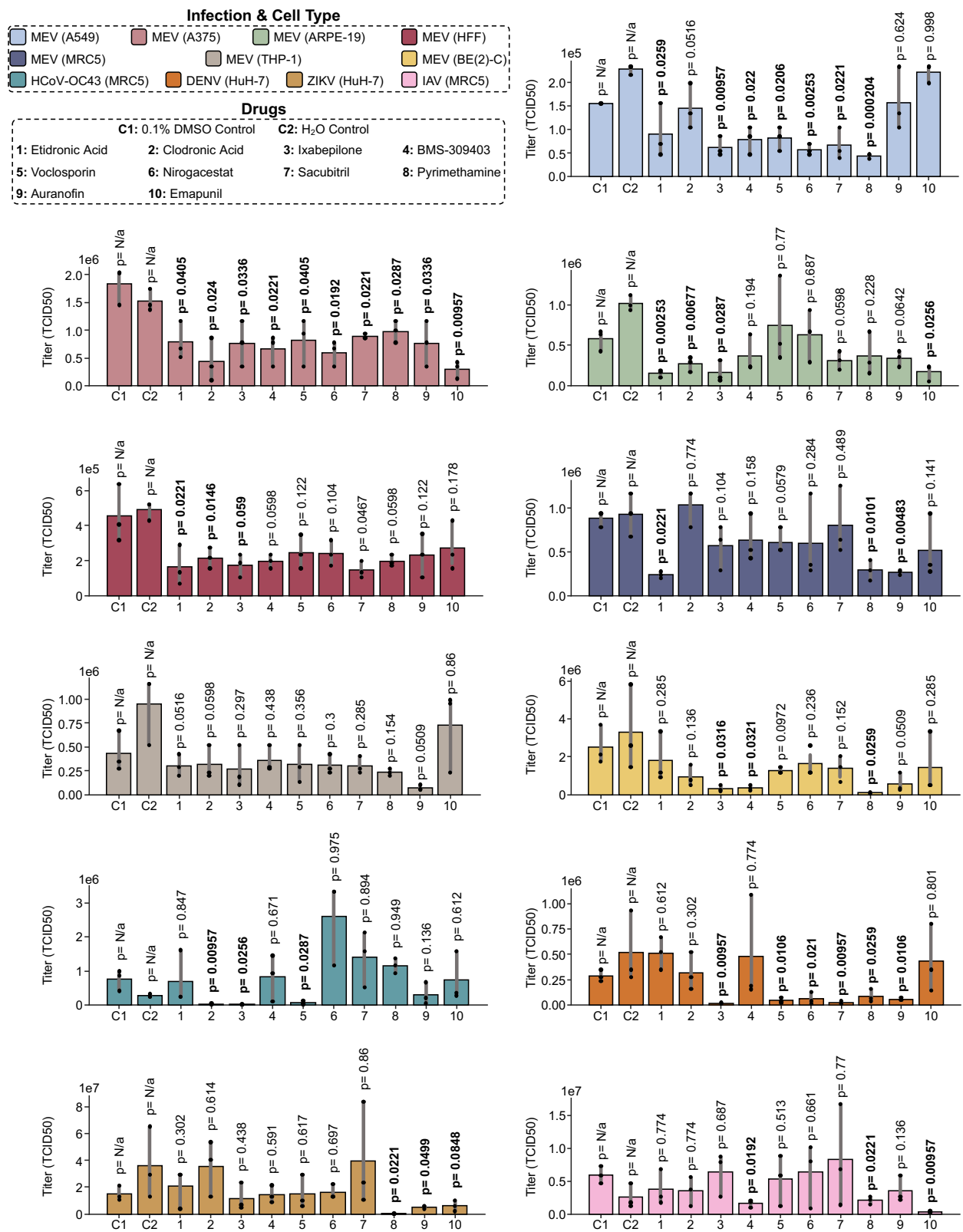

the individual values, and the grey error bar depicts the 95% confidence interval. All experiments were performed in three replicates. P-values (excluding controls) for all samples are listed above each bar and were computed using a one-sided student's T-test comparing the drug treated sample to its appropriate control (either 0.1% DMSO or water treated samples, see methods) P-values were adjusted for multiple comparisons using the Benjamini-Hochberg procedure, and the p-values displayed are the adjusted values.

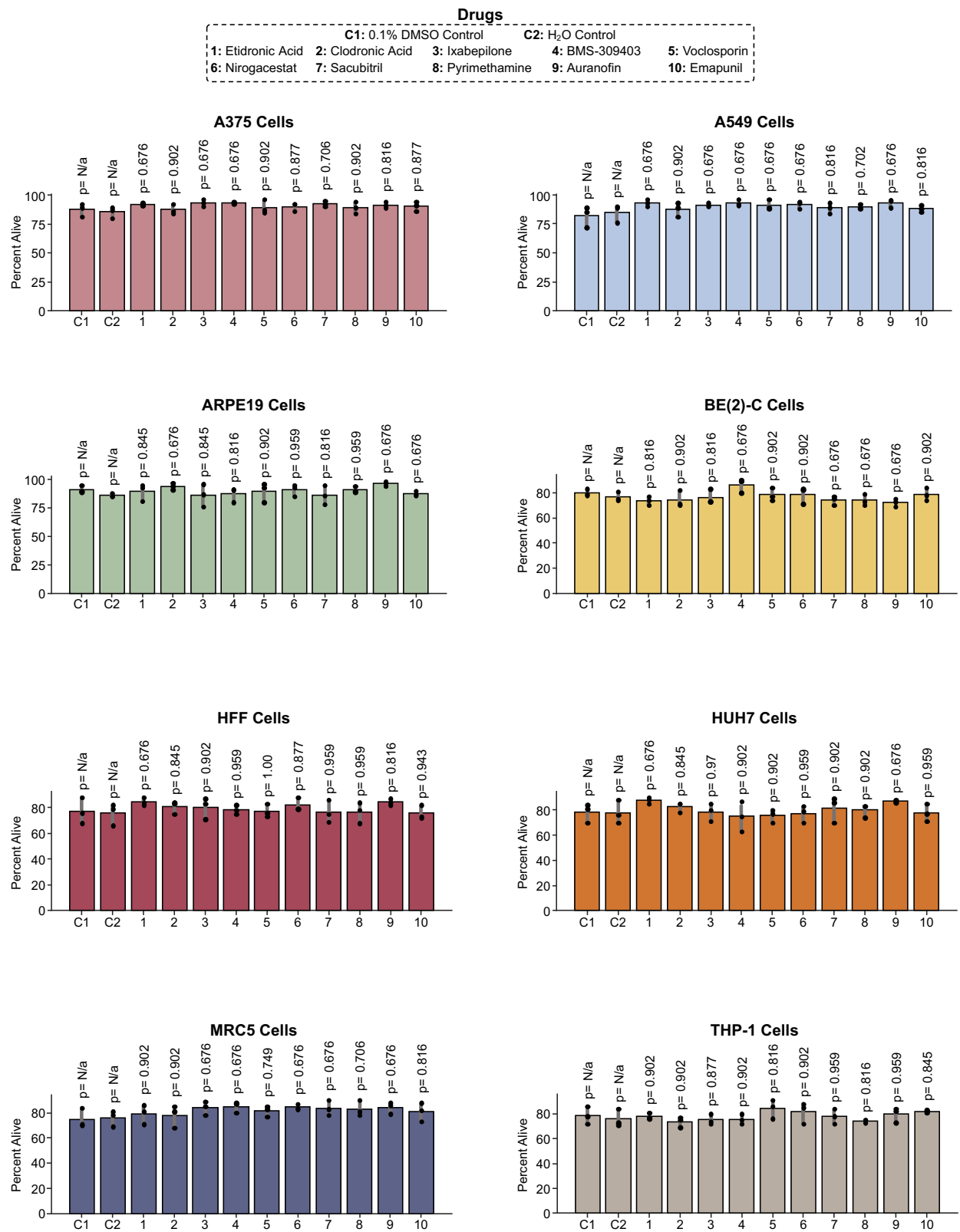

**Figure S12. Uninfected drug treatment control experiments.** Results of trypan blue exclusion assay showing the percentage of alive cells after drug treatment without any viral infections (see

methods). In the plots, the height of the bar is the mean value, and the dots represent the individual values, and the grey error bar depicts the 95% confidence interval. All experiments were performed in three replicates. P-values (excluding controls) for all samples are listed above each bar and were computed using a two-sided student's T-test comparing the drug treated sample to its appropriate control (either 0.1% DMSO or water treated samples, see methods). P-values were adjusted for multiple comparisons using the Benjamini-Hochberg procedure, and the p-values displayed are the adjusted values.

Interactome Regulation Identifies DENV & MeV Infection Similarity Across Cell Lines
(Individual MeV Cell Line Infection Resolved)

■ HCoV-OC43 (MRC5) ■ IAV (MRC5) ■ DENV (HuH7) ■ ZIKV (HuH7)

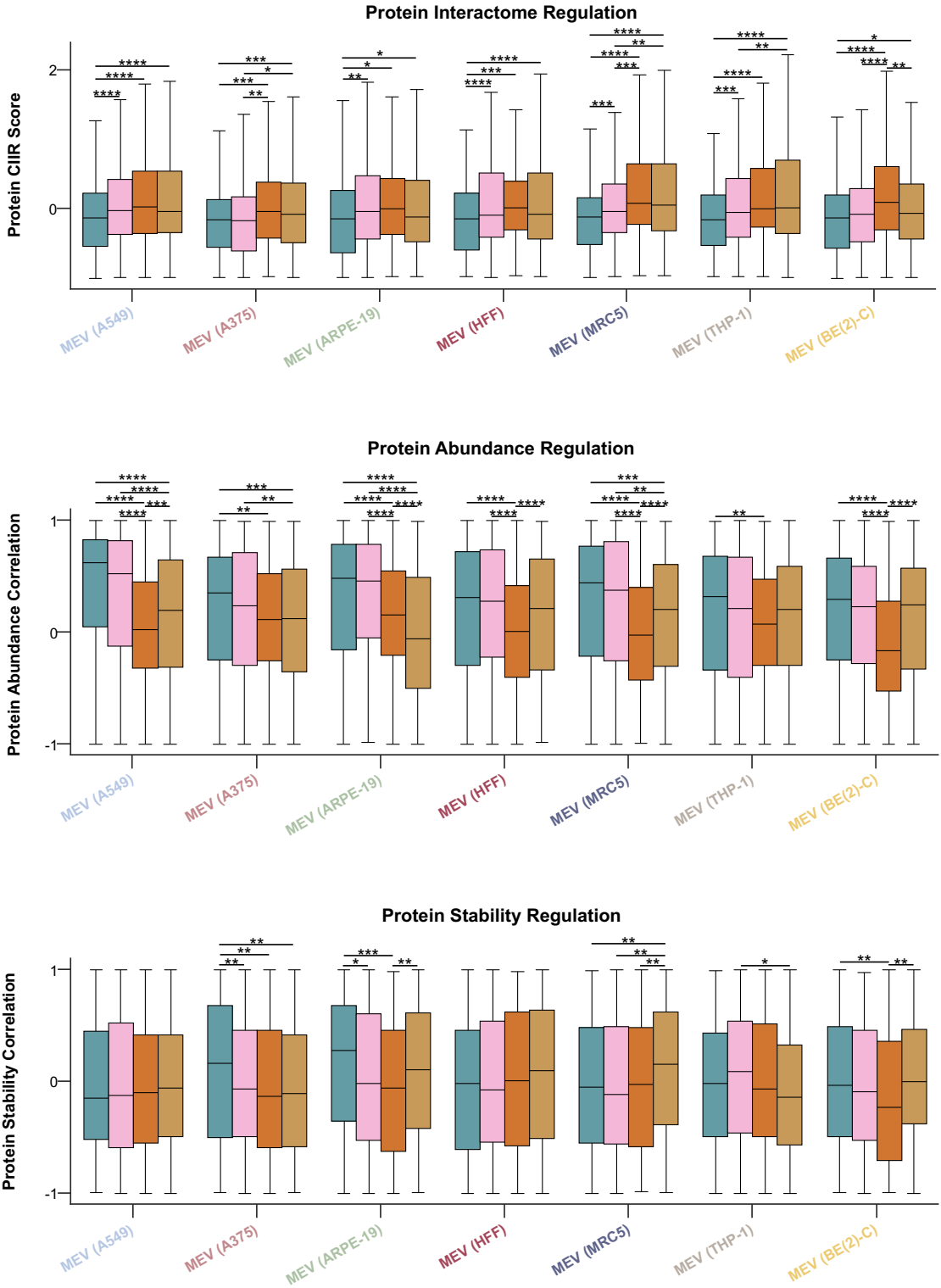

**Figure S13. Comparing convergence of interactome, protein abundance, and protein stability between MeV infections and HCoV-OC43, IAV, DENV and ZIKV infections.** Box plots showing the CIIR score (*top*), protein abundance correlation (*middle*), or protein stability correlation (*bottom*) with MeV infections (resolved by each MeV infection) of top regulated proteins by IIIR score, protein abundance, or thermal stability, respectively, in HCoV-OC43, IAV, DENV, and ZIKV infections. The line within the box represents the median value and the whiskers represent the $\pm 1.5$ interquartile range. P-values were calculated using a two-sided student's T-test and were adjusted for multiple comparisons using the Benjamini-Hochberg procedure. Only p-values $\leq 0.05$ are shown and are represented as \* $\leq 0.05$, \*\* $\leq 0.01$, \*\*\* $\leq 0.001$, and \*\*\*\* $\leq 0.0001$.

#### Temporal Abundance of SHC1 Across MeV Infections

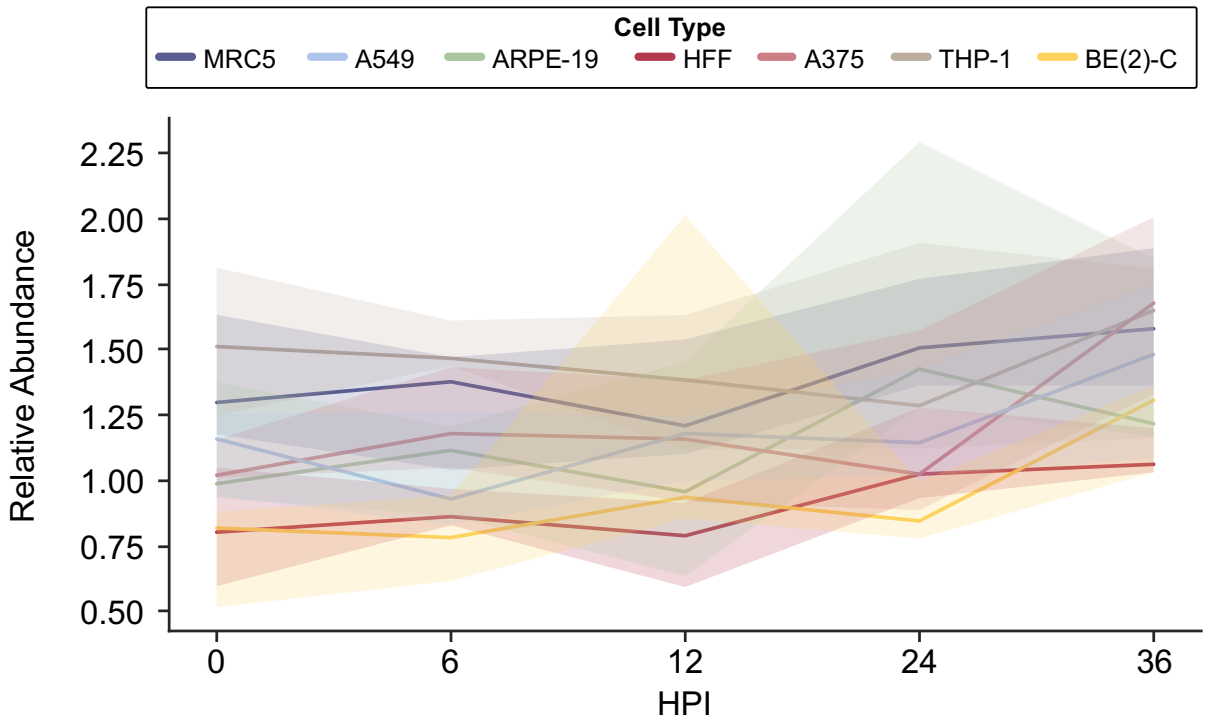

**Figure S14. Temporal SHC1 relative abundance across MeV Infections.** The data has been median normalized, but not normalized to 0 HPI, showing the differences in relative abundance of SHC1 across infections and time. In the plot, the solid line represents the median value, and the shaded region represents the 95% confidence interval.
